# Intracellular pH regulates β-catenin with low pHi increasing adhesion and signaling functions

**DOI:** 10.1101/2024.03.22.586349

**Authors:** Brandon J. Czowski, Angelina N. Marchi, Katharine A. White

## Abstract

Intracellular pH (pHi) dynamics are linked to cell proliferation, migration, and differentiation. The adherens junction (AJ) and signaling protein β-catenin has decreased abundance at high pHi due to increased proteasomal-mediated degradation. However, the effects of low pHi on β-catenin abundance and function have not been characterized. Here, we use population-level and single-cell assays to show that low pHi stabilizes β-catenin, increasing junctional, cytoplasmic, and nuclear abundance. We assayed single-cell protein degradation rates to show that β-catenin half-life is longer at low pHi and shorter at high pHi compared to control. Importantly, a constitutively stabilized and pHi-insensitive β-catenin mutant (β-catenin-H36R), has a longer and pHi-independent half-life. We also determined that the pH-dependent stability of β-catenin affects both its adhesion and signaling functions. We show that the composition of AJs changes with pHi; at low pHi, E-cadherin-containing AJs are enriched in β-catenin while plakoglobin abundance is reduced. Conversely, when β-catenin is lost from E-cadherin-containing AJs at high pHi, plakoglobin is increased. We also found that cell area was reduced at low pHi and increased at high pHi compared to control while cell volume was unaffected, suggesting pHi alters cell-cell adhesion. Finally, we show that low pHi increases β-catenin transcriptional activity in single cells and is indistinguishable from a Wnt-on state, while high pHi reduces β-catenin transcriptional activity compared to control cells. This work characterizes pHi as a true rheostat regulating β-catenin abundance, stability, and function, solidifying β-catenin as a molecular mediator of pHi-dependent cell processes via pH-dependent adhesion and signaling functions.

**Summary:** Intracellular pH (pHi) regulates the degradation rate of the pH sensor β-catenin, altering protein abundance, subcellular localization, and function in epithelial cells. This work shows pHi acts as a rheostat to alter both adhesion and signaling functions of β-catenin.

## Introduction

Spatiotemporal pHi dynamics regulate cell processes such as differentiation (1), cell migration(2), cell cycle progression (3), and cell fate determination (4). While low pHi is required to maintain the adult stem cell niche (1, 4) and transient increases in pHi are required for successful differentiation (1), the molecular drivers of these pHi-dependent processes are unknown. Recent work identified β-catenin as a pH-sensitive protein with decreased abundance at high pHi driven by pH-dependent binding of β-catenin to the E3 ligase, β-transducin repeat-containing protein (β-TrCP) (5).

β-catenin is a multifunctional protein with structural and transcriptional roles, maintaining adherens junctions (AJs) between epithelial cells and functioning as the main signal transducer of the Wnt pathway (6–8). Co-localization of β-catenin with E-cadherin at AJs preserves junction integrity (9), protects E-cadherin from degradation (10), and recruits α-catenin to link AJs to the cytoskeleton (11). In the absence of Wnt ligand, cytoplasmic β-catenin is phosphorylated by casein kinase 1 α (CK1α) and glycogen synthase kinase 3-β (GSK3β), enabling binding and ubiquitination by β-TrCP (12) and subsequent proteasomal degradation (13). Wnt activation inhibits the destruction complex (14): kinase activity is reduced and β-TrCP is dissociated (15, 16). Non-phosphorylated (active) β-catenin accumulates in the cytoplasm, translocates to the nucleus, and activates transcription of proliferative and morphogenic genes (17) (18) (19) by displacing Groucho in the repressive Groucho/TCF/LEF complex (20). The dual roles of β-catenin in AJs and Wnt signaling are essential for cell differentiation, stem cell maintenance, and tissue homeostasis (21, 22). Thus, the 2018 paper showing that high pHi decreases β-catenin stability (5) led several groups to subsequently suggest pH-dependent stabilization of β-catenin may underlie pH-dependent cell differentiation and tissue morphogenesis (4, 23).

However, while White et al. previously showed that high pHi reduces whole-cell and junctional β-catenin abundance in the *Drosophila* eye epithelia and in mammalian epithelial cells (5), the work had several significant limitations. First, the effect of decreased pHi on β-catenin abundance was not examined due to the lack of reliable methods for lowering pHi in epithelial cells. Second, the impact of pHi-mediated β-catenin abundance changes on AJ integrity and composition was not fully characterized. Third, the pH-dependent transcriptional activity of endogenous β-catenin was not investigated. Given the established context-dependent function of β-catenin, our understanding of how changes in pHi regulate β-catenin abundance and function remains incomplete. Thus, a full characterization of the effects of both high and low pHi on the adhesion and signaling roles of β-catenin is needed.

Here, we determined pH-dependent effects on β-catenin abundance, subcellular localization, degradation dynamics, and function using methods to both raise and lower pHi in Madin-Darby Canine Kidney cells. Using population-level immunoblots, we demonstrate pH-dependent β-catenin abundance, with increased abundance at low pHi compared to high pHi. We also used immunofluorescent microscopy to characterize the effects of pHi on the abundance of distinct subcellular pools of β-catenin within single cells. We show that nuclear, cytoplasmic, and junctional β-catenin pools are all decreased at high pHi while low pHi stabilizes both the nuclear and cytoplasmic pools of β-catenin. We also determined β-catenin degradation dynamics in single cells using a photoconvertible mMaple3-β-catenin fusion. We show that low pHi increases β-catenin half-life while high pHi reduces its half-life. Furthermore, a constitutively stabilized β-catenin mutant (β-catenin H36R) where the pH-sensitive His36 is mutated to a non-titratable arginine, exhibits a high and pH-insensitive half-life. Collectively, our data demonstrate that pHi functions as a rheostat to regulate β-catenin degradation rates.

Importantly, our work also characterizes how pHi regulates β-catenin function via altering AJ composition and transcriptional activation. At low pHi, β-catenin is enriched in E-cadherin-containing AJs, while high pHi depletes β-catenin from E-cadherin-containing AJs. However, recruitment of plakoglobin at high pHi maintains AJ integrity and epithelial morphology. We also found pH-dependent changes in cell shape but not volume, a phenotype normally associated with altered cell adhesion. We found that low pHi reduces cell cross-sectional area and high pHi increases cell cross-sectional area compared to control. Additionally, we found that modest (∼10%) increases in nuclear abundance of total β-catenin and active (non-phosphorylated) β-catenin at low pHi significantly increases β-catenin transcriptional activity to levels indistinguishable from those observed with addition of extracellular Wnt ligand. Our findings establish that pHi functions as a rheostat to regulate the stability and function of endogenous wild-type β-catenin. This work improves our understanding of the functional biochemical outcomes of pH-dependent β-catenin regulation that may underlie both normal (differentiation) and pathological (transformation, cancer stem cells) processes marked by temporal changes in pHi (1, 24, 25).

## Results

### Intracellular pH differentially affects subcellular pools of β-catenin

To investigate the pH-dependent regulation of β-catenin in a mammalian cell model, we used Madin-Darby canine kidney (MDCK) cells. β-catenin has been extensively studied in MDCK cells, where it was previously shown that increased pHi was sufficient to reduce overall β-catenin abundance at the population level as well as nuclear and junctional pools at the single-cell level(5). This prior work established a role for high pHi in regulating β-catenin abundance; however, the lack of methods to lower pHi in this cell line prevented complete characterization of the role of pHi in regulating β-catenin abundance, localization, and function in epithelial cells.

To experimentally lower pHi in MDCK cells, we used the combination of a sodium-proton exchanger 1 (NHE1) inhibitor (5-(N-ethyl-N-isopropyl)amiloride (EIPA)) and a sodium-bicarbonate cotransporter (NCBn1) inhibitor (2-chloro-N-[[2′-[(cyanoamino)sulfonyl][1,1′-biphenyl]-4-yl]methyl]-N-[(4-methylphenyl)methyl]-benzamide (S0859)) (26). To experimentally raise pHi, we used 15% CO_2_ atmospheric culture conditions, which increases the bicarbonate concentration in the medium and, through the function of NCBn1, increases intracellular bicarbonate to raise pHi (5). Basal pHi in control MDCK cells was 7.36±0.02, while pHi manipulation treatments significantly increased pHi to 7.49±0.10 (15% CO_2_) and significantly decreased pHi to 7.17±0.05 (EIPA+S0859) after 24 hours (**Figure 1A**) with no change in cell viability (**Figure S1A**). Importantly, treatment with proteosome inhibitor MG132 (27) did not affect the magnitude of pHi changes achieved with these protocols (**Figure 1A**). Together, these manipulation methods enable us to study how physiological changes in pHi of 0.1-0.3 pH units regulate β-catenin abundance and function. These pHi changes are physiologically relevant as pHi dynamics of similar magnitude occur during cellular differentiation (+0.2 pH units) (1) and cell division (±0.25 pH units) (3).

**Figure 1:**
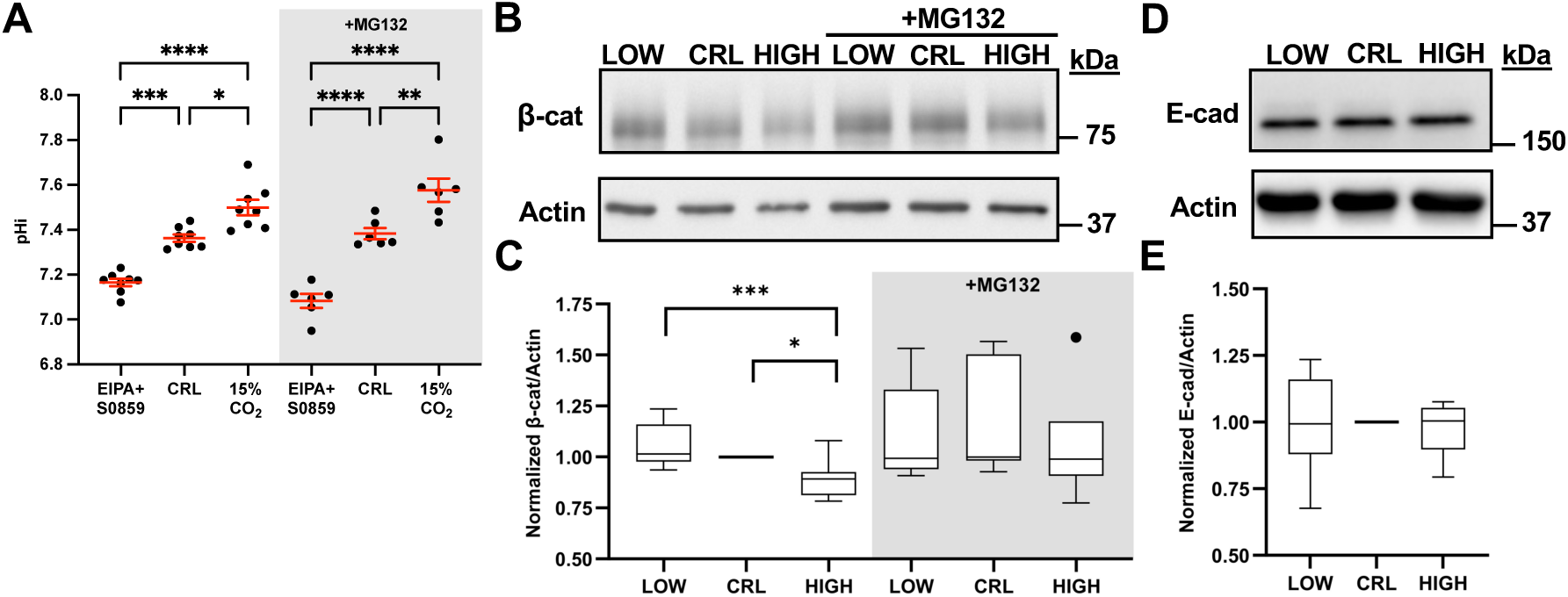
Decreased pHi stabilizes β-catenin abundance through a proteosome-dependent mechanism. **(A)** Box and whisker plots showing population level pHi measurements in MDCK cells after 24-hour treatments with 1 µM EIPA+ 30 µM S0859 (low) or 15% CO_2_ (high) to lower and raise pHi, respectively. Mean and SEM shown. n=6-8. **(B)** Representative immunoblot of β-catenin (β-cat) and actin under low, control, and high pHi conditions in the presence and absence of proteosome inhibition (MG132). **(C)** Quantification of β-catenin immunoblot data collected as described in B. Individual biological replicates were normalized to control MDCK within each experiment. Box and whisker plots show median (line), 25^th^-75^th^ percentile (boxes), min and max (whiskers), and outlier values with points. n=6-8. **(D)** Representative immunoblot of E-cadherin and actin under low, control, and high pHi conditions. **(E)** Quantification of E-cadherin immunoblot data collected as described in D and displayed and normalized as described in C. n=6. Statistical analyses in A were determined using one-way ANOVA test with Sidak’s correction for multiple comparisons. Statistical analyses in C and E were determined with ratio paired t-tests between treatment groups and one-sample t-tests with a hypothetical mean of 1.0 when comparing to control (which had no variation due to normalization). *P<0.05; **P<0.01 ***P<0.001; ****P<0.0001.

We first used western blot approaches to assay β-catenin abundance at high and low pHi. We confirmed that high pHi decreased β-catenin abundance compared to control and found that low pHi was sufficient to stabilize β-catenin abundance compared to high pHi (**Figure 1 B, C**). Importantly, we demonstrated that pH-dependent abundance of β-catenin is regulated by proteosome-mediated degradation, as treatment with proteasome inhibitor (MG132) abrogated pH-dependent abundance (**Figure 1B, C**). To ensure that low pHi was not inducing global protein stabilization, we probed the abundance of the AJ protein, E-cadherin (**Figure 1D**). We found that high and low pHi conditions had no effect on the overall abundance of E-cadherin (**Figure 1E**) or actin (**Figure S1B**), demonstrating that the observed pH-dependent changes in β-catenin abundance are not driven by global protein degradation or stabilization. Previous work showed that phosphorylation of β-catenin was unchanged with increased pHi (5), we similarly found that phospho-β-catenin levels were not significantly different with altered pHi (**Figure S2A**), demonstrating that protein abundance differences are not a result of pH-sensitive phosphorylation of β-catenin’s degron motif. Taken together, these data suggest that pHi functions as a rheostat in regulating β-catenin abundance at the population level through a pHi- and proteasome-dependent mechanism, with increased stability at low pHi compared to high pHi.

Because the function of β-catenin is in part determined by its subcellular localization and availability of binding partners (28–30), we next investigated how pHi alters the abundance and distribution of each subcellular pool of β-catenin. We first confirmed that the pHi manipulation techniques altered pHi in single MDCK cells using live-cell, confocal microscopy with nigericin standardization (3, 31) (**Figure S3A**). We found that under control conditions, the median pHi value of MDCK cells was 7.39±0.14 (**Figure S3B**), in line with our previous calculations using population-level measurements (**Figure 1A**). Importantly, treatment with EIPA+S0859 significantly lowered single-cell pHi (7.26±0.17), while treatment with 15% CO_2_ significantly raised single-cell pHi (7.42±0.20), relative to control MDCK cells (**Figure S3B**).

We hypothesized that high pHi would decrease β-catenin levels in each subcellular pool, whereas low pHi would stabilize each subcellular pool. This hypothesis is predicated on recent data suggesting subcellular pools of β-catenin can rapidly exchange and that exchange is regulated by the availability of subcellular binding partners (28, 29). To test this hypothesis, we developed a 3D image analysis pipeline to segment normal epithelial cells (MDCK) and separately measure β-catenin abundance in the nucleus, cytoplasm, and at adherens junctions (Figure S3, see methods). Briefly, cytoplasmic and nuclear regions of interest (ROI) were generated in Nikon Imaging Software-Elements (NIS-Elements), using Hoechst dye (DNA) to identify nuclear boundaries and an E-cadherin antibody (adherens junctions) to identify cell boundaries (**Figure S4**). This segmentation was performed for every cell in the field of view, where the pixel intensity within cytoplasmic or nuclear regions of interest were averaged and exported for comparison. To specifically quantify junctional pools of β-catenin, we used IMARIS (Oxford Instruments, see methods) software to generate 3D surfaces using E-cadherin intensity as a marker of adherens junctions. This analysis pipeline enables voxel-by-voxel quantification within each E-cadherin-containing AJ surface.

We first assessed the staining patterns of β-catenin and E-cadherin in MDCK cells (**Figure 2A**). Importantly, the subcellular distribution of β-catenin in MDCK cells agrees with previous reports, with abundant membrane localization of β-catenin, weak staining of the cytoplasm, and very weak staining of nuclear β-catenin as expected for epithelial cells that are in a Wnt-inactive state (14). Using the image segmentation procedure described above, we quantified β-catenin abundance in each subcellular pool (cytoplasmic, nuclear, and junctional) after 24 hours of pHi manipulation (**Figure 2A**). We found that all subcellular pools of β-catenin decreased at high pHi (**Figure 2B**), which agrees with the population-level western blot data, and our hypothesis of increased pHi being sufficient to reduce β-catenin abundance across all subcellular pools. This data is also in agreement with prior quantification of β-catenin abundance at junctions and in the nucleus of MDCK cells maintained at high pHi using an alternative method (ammonium chloride) (5). At low pHi, we found that the cytoplasmic and nuclear pools of β-catenin were significantly increased compared to both control and high pHi conditions (**Figure 2A, B**). In our initial analysis, low pHi was not sufficient to significantly increase the abundance of β-catenin at E-cadherin-containing AJ compared to control MDCK cells (**Figure 2B**).

**Figure 2:**
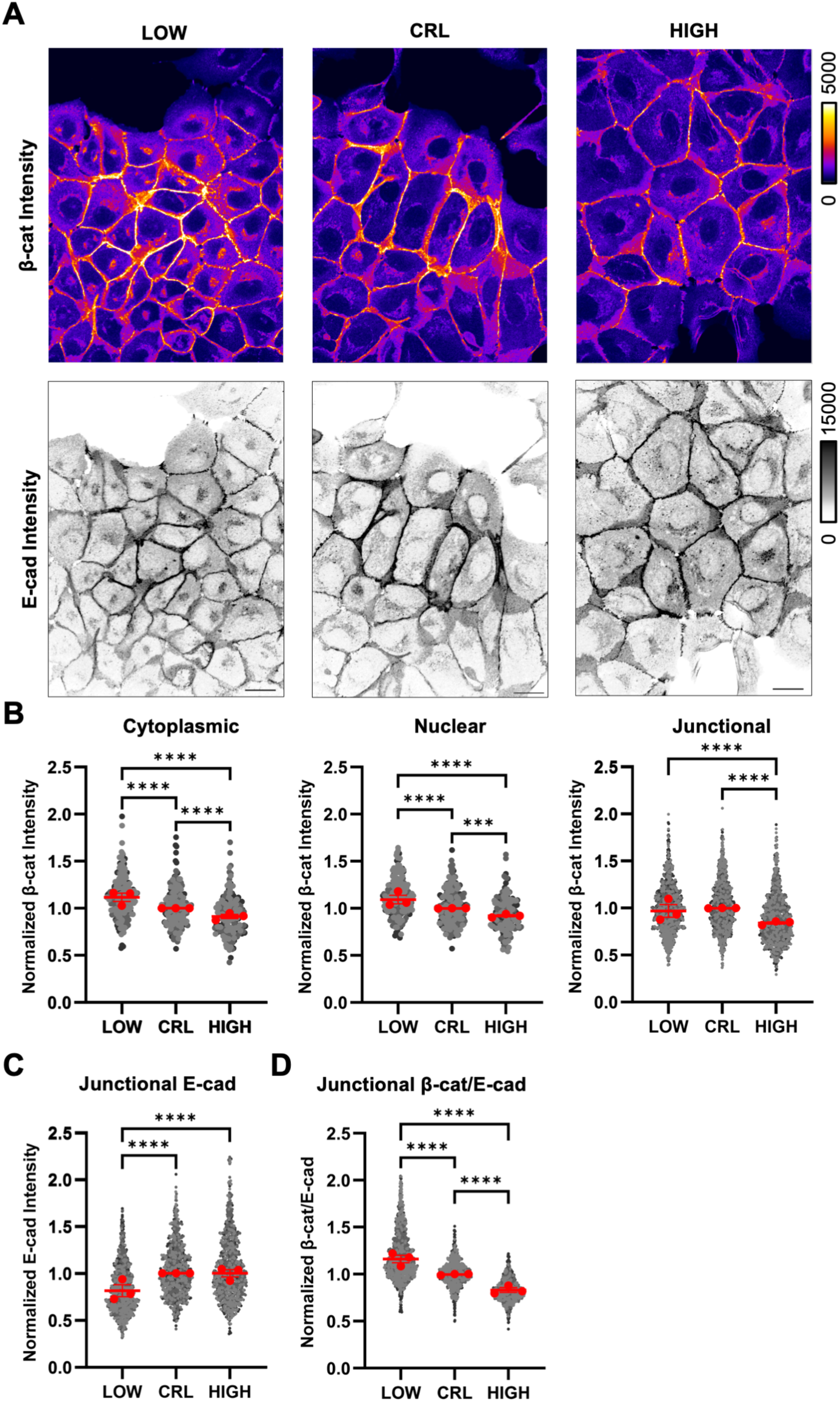
β-catenin abundance across subcellular pools is differentially regulated by pHi. **(A)** Representative confocal images of MDCK cells fixed and stained for β-catenin (β-cat) and E-cadherin (E-cad). β-catenin is pseudocolored and E-cadherin is shown in inverse mono according to each scale. Scale bar: 20 µM. **(B)** Quantification of cytoplasmic, nuclear, and junctional pools of β-catenin. Individual cells and surfaces were normalized to the median of control conditions within each biological replicate. Individual cytoplasms, nuclei, and junctions are shown as gray dots with various shading for each biological replicate. Red dots represent the median for each biological replicate with interquartile ranges shown with bars. **(C)** Quantification of junctional E-cadherin displayed and normalized as in B. **(D)** Calculation of β-cat/E-cad ratio from quantifications in B and C that are normalized and displayed as in B. Individual cytoplasms and nuclei, n=171-221; junctions, n=934-1194, from 3 biological replicates. Statistical analyses in B-D were performed using the Kruskal-Wallis test with Dunn’s multiple comparison correction. ***P<0.001, ****P<0.0001.

To ensure we were not missing an effect, we quantified E-cadherin intensity within the AJ surfaces. Contrary to our population level measurements of total E-cadherin (**Figure 1C**), we observed a significant decrease in E-cadherin at MDCK junctions under low pHi compared to control MDCK (**Figure 2A, C**). To control for this loss of E-cad at low pHi, we next quantified the ratio of β-catenin to E-cadherin within single MDCK junctions generated using IMARIS. We observed strong pH-dependence of β-catenin/E-cad ratios with increased β-catenin in E-cadherin-containing AJs at low pHi compared to control and high pHi (**Figure 2D**). This suggests that the level of β-catenin saturation at AJs changes with pHi, with less β-catenin in E-cadherin-containing AJs at high pHi and more β-catenin in E-cadherin-containing AJs at low pHi, even though total junctional E-cadherin is reduced at low pHi.

Quantification of each pool of β-catenin provides information as to how β-catenin subcellular distribution and abundance change with respect to both increased and decreased pHi. Taken together, our data suggest that subcellular pools of β-catenin are exchangeable, with high pHi reducing cytoplasmic, nuclear, and junctional β-catenin. Similarly, our data suggest low pHi results in cytoplasmic and nuclear accumulation and a relative increase in β-catenin at E-cadherin-containing AJs.

As we were analyzing these data, we noted a distinct change in cell morphology of the fixed cells across our treatment conditions, with apparent pH-dependent cell size changes (**Figure 2A**). To quantify these changes in live cells, we used CellMask DeepRed to label the plasma membranes of live MDCK cells under low, control, and high pHi conditions and collected 3D confocal image stacks of the cells to quantify both cell cross-sectional area and volume (**Figure S5A**). While cell volume was unchanged across pHi treatment conditions (**Figure S5B**), cell cross-sectional area decreased at low pHi compared to control MDCK and increased at high pHi compared to control (**Figure S5C**). Prior work has shown that cell cross-sectional area increases when cells have higher cell-matrix adhesion, while cell cross-sectional area is smaller in cells with high cell-cell adhesion (32). Thus, our data demonstrate that pH-dependent changes in junctional β-catenin can drive cell shape changes that are associated with altered cell-cell adhesion. We next probed whether AJ composition was altered by pHi dynamics.

### Endogenous plakoglobin compensates for pHi-dependent loss of β-catenin to maintain AJs at high pHi

In the immunofluorescence assays of β-catenin, we observed no change in MDCK colony formation with altered pHi, even when junctional pools of β-catenin were significantly decreased at high pHi (**Figure 2A, B**) and cell cross-sectional area was increased (**Figure S5**). This suggests that other AJ proteins may be able to rescue the loss of junctional β-catenin to maintain AJ integrity. While β-catenin plays a crucial role in maintaining AJs in epithelial cells, plakoglobin serves adhesive functions in both AJs and desmosomes, with preferential incorporation into the latter (33). Work by Fukunaga and colleagues demonstrated the functional redundancy of β-catenin and plakoglobin in maintaining AJ integrity, where cell-cell contacts can be rescued with plakoglobin expression in a β-catenin null background (34). Furthermore, knockout of β-catenin in mouse models resulted in significant recruitment of plakoglobin to AJs, compensating for the loss of AJ-associated β-catenin (35). A similar effect was observed in mouse models using conditional β-catenin knockdown in cardiomyocytes, where β-catenin was reduced by 90% and plakoglobin was subsequently enriched at junctions (36). We therefore hypothesized that in MDCK cells, the homologous protein plakoglobin can rescue β-catenin loss from AJs at high pHi to preserve MDCK epithelial morphology.

To determine whether plakoglobin compensates for the loss of β-catenin at AJs at high pHi, we performed immunofluorescent confocal microscopy of plakoglobin using the same segmentation pipeline (outlined in **Figure S4**) after 24 hours of pHi manipulation. We found that junctional-associated plakoglobin significantly increased at high pHi and significantly decreased at low pHi compared to control MDCK (**Figure 3A, B & C**). Importantly, overall abundance of plakoglobin is unchanged by pHi (**Figure 3D & E**) suggesting a change in localization is driving these observations. These data, combined with the analysis of β-catenin abundance at E-cadherin-containing junctions (**Figure 2**), do suggest that pHi dynamics can remodel AJ composition. However, the ratio-ed voxel-by-voxel analysis are still limited as we are comparing median ratioed intensity values that are then normalized from one replicate to the next. We wanted to determine functional correlations between expression level of one AJ protein and another in our data.

**Figure 3.**
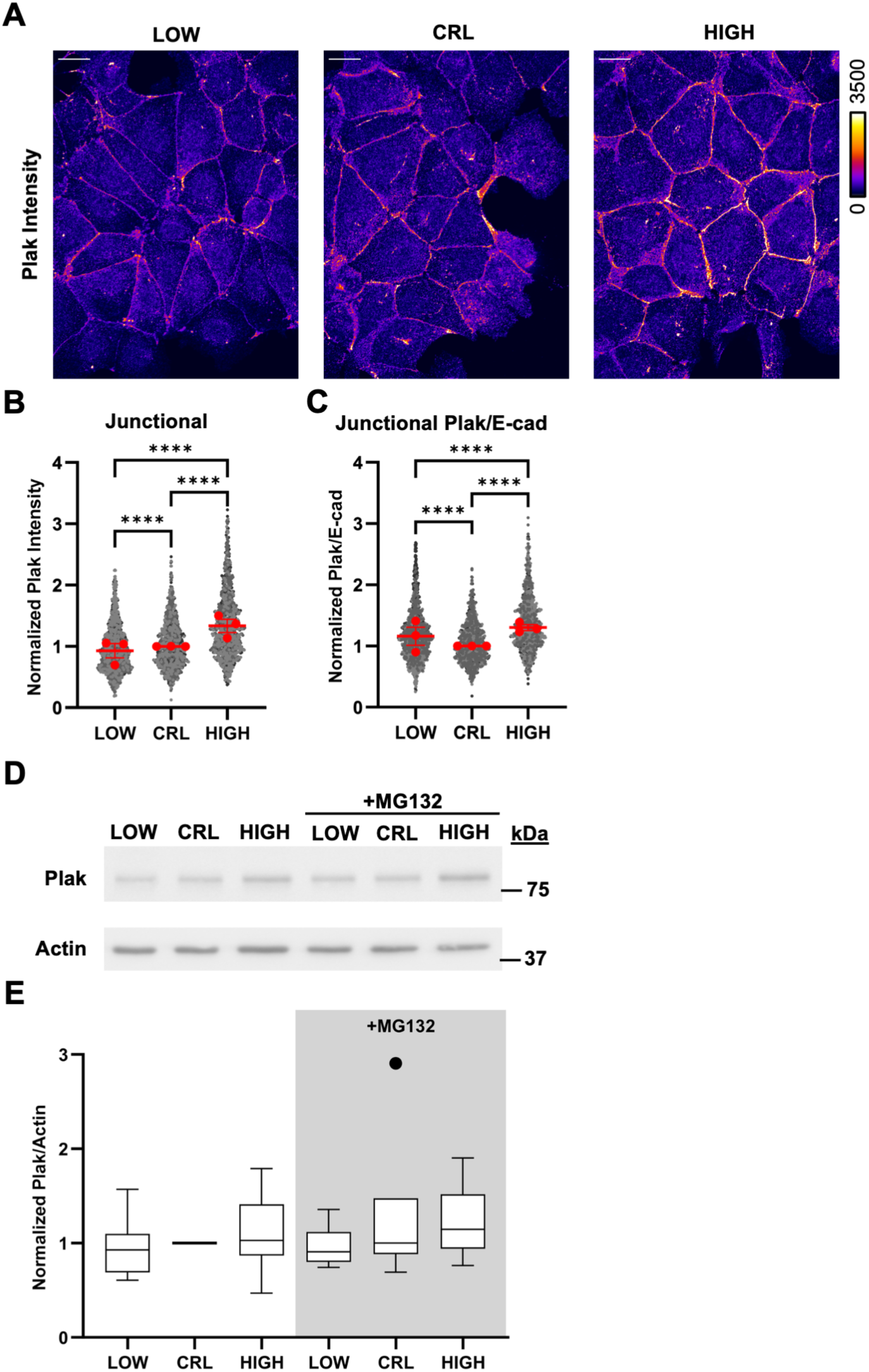
Endogenous plakoglobin compensates for loss of β-catenin from junctions at high pHi. **(A)** Representative confocal images of MDCK cells fixed and stained for Plakoglobin (Plak). Plakoglobin is pseudocolored according to scale. Scale bar: 20 µM. Quantification of junctional pools of Plakoglobin **(B)** and Plak/E-cad ratios **(C)** from individual surfaces. Individual surfaces were normalized to the median of control conditions within each biological replicate. Individual junctions are shown as gray dots with various shading for each biological replicate, n=934-1194, from 3 biological replicates. Red dots represent the median for each biological replicate with interquartile ranges shown with bars. **(D)** Representative immunoblot of Plak and actin under low, control, and high pHi conditions in the presence and absence of proteosome inhibition (MG132). **(E)** Quantification of Plak immunoblot data collected as described in D. Individual biological replicates were normalized to control MDCK within each experiment. Box and whisker plots show median (line), 25^th^-75^th^ percentile (boxes), min and max (whiskers), and outlier values with points; from 5-6 biological replicates. Statistical analyses of B and D were performed using the Kruskal-Wallis test with Dunn’s multiple comparison correction. Statistical analyses of C and E were determined with ratio paired t-tests between treatment groups and one-sample t-tests with a hypothetical mean of 1.0 when comparing to control (which had no variation due to normalization). ****P<0.0001.

Thus, to characterize AJ composition in more detail, we generated voxel-by-voxel correlation plots of normalized β-catenin and plakoglobin intensities vs E-cadherin intensities in E-cadherin containing AJs. Importantly, because we performed simultaneous labeling of E-cadherin, plakoglobin, and β-catenin, these correlation maps are robust across biological replicates. As expected, we found that both β-catenin and plakoglobin intensities were positively correlated with E-cadherin intensity (**Figure S6A, B**). When we analyzed the correlation plots of β-catenin, we found a significant increase in β-catenin abundance at low pHi compared to high pHi, as evidenced by the distinct separation of scatter plot data by pHi ((shifted along the y axis (β-catenin intensity)) (**Figure S6A**). When we analyzed the correlation plots of plakoglobin vs. E-cadherin, we found no change in overall plakoglobin abundance, as evidenced by the overlap of scatter plot data by pHi (**Figure S6B**).

Taken together, these correlation data recapitulate the pH-dependent abundance data showing β-catenin has pHi-dependent abundance (**Figure 2**) while plakoglobin does not (**Figure 4**). These scatter plots additionally show that low pHi uniformly increases β-catenin abundance in E-cadherin-containing voxels independently of E-cadherin concentration. In support of this, we found correlation coefficients between E-cadherin and β-catenin were unchanged with changing pHi (**Figure S6C**). This suggests that the observed β-catenin abundance changes at AJs (**Fig 2 and Fig. S6A**) are not being driven by changes in β-catenin affinity for E-cadherin (which would be dependent on the concentration of E-cadherin and thus affect correlations) but instead are driven by overall changes in β-catenin abundance (which would increase binding independently of E-cadherin concentration). Importantly, we found that β-catenin had higher correlation with E-cadherin at low pHi (Spearman r=0.71) compared to plakoglobin (Spearman r=0.43, p<0.0001) (**Fig. S6C,D**). We also found that plakoglobin was more highly correlated with E-cadherin at high pHi (Spearman r=0.55) compared to low pHi (Spearman r=0.43; p<0.0001) (**Fig. S6C,D**), suggesting plakoglobin abundance is not being directly regulated and instead it is being re-localized to AJs at high pHi in a concentration-dependent manner.

**Figure 4.**
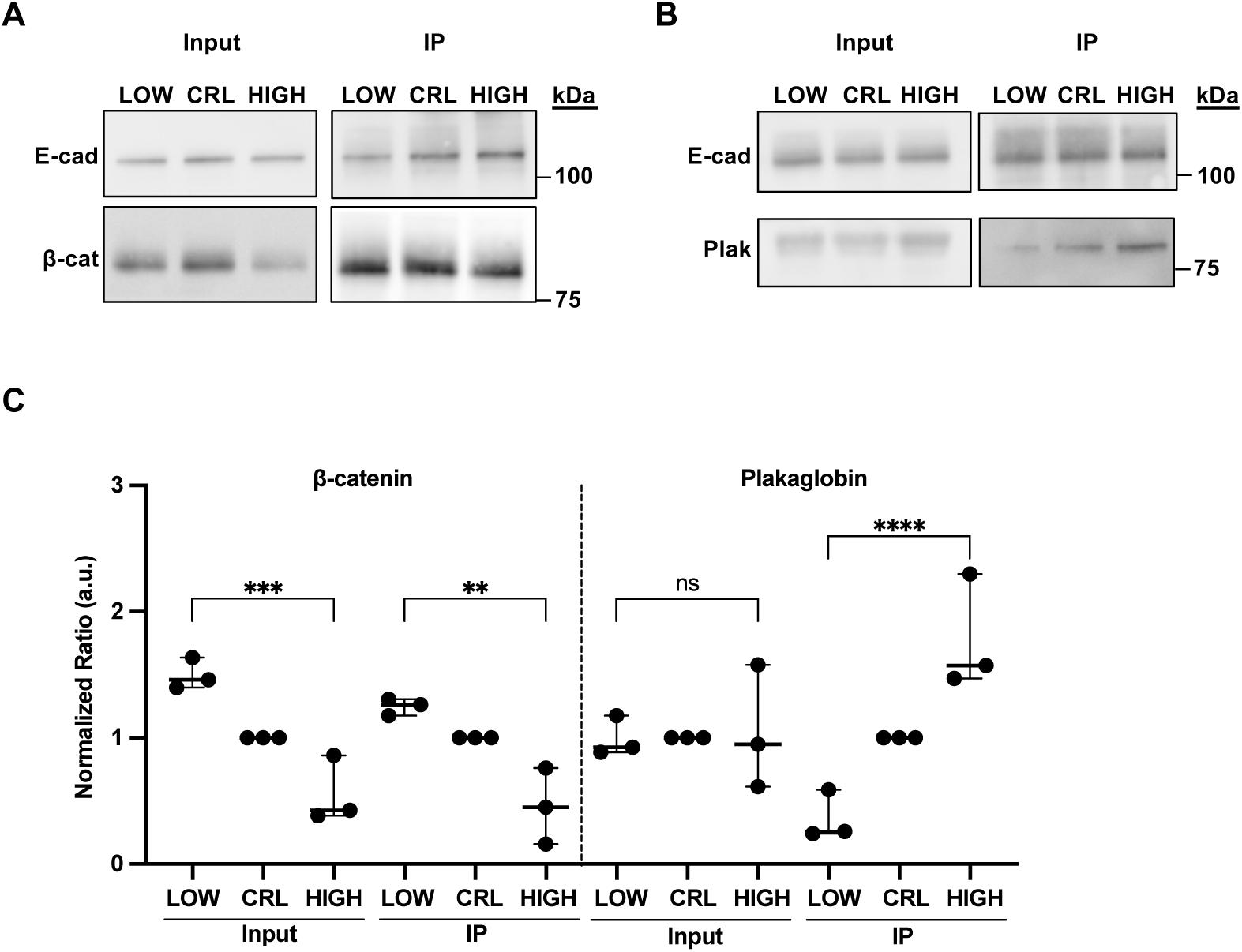
Immunoprecipitation of E-cadherin shows AJ composition is altered with changes in pHi. **(A)** Representative immunoblots of plakoglobin (Plak) and E-Cadherin in whole cell lysates (left, input) and in E-cadherin containing immune complexes (right, IP). **(B)** Representative immunoblots of β-catenin (β-cat) and E-Cadherin in whole cell lysates (left, input) and in E-cadherin containing immune complexes (right, IP). **(C)** Quantification of plakoglobin and β-catenin in E-cadherin immunocomplexes collected as described in A. Individual biological replicates were normalized to control MDCK within each experiment. Scatter plots show all data points from n=3 biological replicates. Statistical significance was determined using a one-way ANOVA between high and low treatment groups. *P<0.05, **P<0.01, ***P<0.001, ****P<0.0001.

We next biochemically confirmed that pHi dynamics alter the composition of E-cadherin-based AJs. We prepared lysates from MDCK cells under low, control, or high pHi conditions and performed immunoprecipitation of E-cadherin. We found that β-catenin was enriched in E-cadherin immune complexes at low pHi compared to control and high pHi (**Figure 4A, C**). Conversely, plakoglobin was enriched in E-cadherin immune complexes at high pHi compared to control and low pHi (**Figure 4B, C**). This result confirms the pH-dependent co-localization and correlation data shown in single cells (**Figures 2, 3, and S6**). In summary, our data show that while β-catenin co-localization with E-cadherin is reduced at high pHi compared to control (**Figures 2D, 4B**), plakoglobin co-localization with E-cadherin is increased (**Figures 3C, 4C**). While further work is required to fully characterize pH-dependent AJ remodeling, our data suggest that when β-catenin is lost from junctions at high pHi, plakoglobin is recruited to stabilize AJs. Similarly, when β-catenin is stabilized at E-cadherin-containing AJs at low pHi, plakoglobin abundance is reduced.

### Characterization of pH-dependent β-catenin degradation

Prior work has already extensively examined roles for high pHi in reducing β-catenin stability (5), but the effects of low pHi on β-catenin stability have not been characterized. We next measured the temporal localization and stability dynamics of endogenous β-catenin at the population level, with the prediction that low pHi increases β-catenin stability resulting in the observed increased β-catenin abundance at low pHi. We first quantified pH-dependent stability of β-catenin by performing a cycloheximide chase assay (37) after 24 hours of pHi manipulation in MDCK cells, collecting samples every 30 minutes for 2.5 hours (**Figure S7A**). The half-life of endogenous β-catenin under control condition was determined to be 1.58 hours, in agreement with prior work in MDCK cells(5). Importantly, we found that β-catenin half-life was increased at low pHi (2.79 hours) and decreased at high pHi (1.22 hours) compared to control (**Figure S7B**). Of note, White et al. used different treatment conditions to raise pHi (ammonium chloride compared to our method of 15% CO_2_) (5). Thus, our work indicates that high pHi accelerates β-catenin degradation, regardless of the method of pHi manipulation. Finally, this work is the first to report an extension of β-catenin half-life in cells under low pHi conditions.

### An experimental approach to monitor β-catenin degradation rates in single cells

The population-level analyses of β-catenin stability using cycloheximide demonstrated pH-dependent stability, where low pHi increased and high pHi decreased β-catenin stability (half-life) relative to control. Although these data revealed altered rates of β-catenin degradation with pHi manipulation, these population-level analyses require discontinuous measurements across cell populations and lack both single-cell and temporal resolution. Single-cell analyses are important to better connect and support emerging roles for pH-dependent β-catenin regulation in processes that occur at the single-cell level, such as differentiation (1). One approach for quantifying live- and single-cell protein dynamics is to monitor a photoconvertible fluorescent protein (PCFP) fused to the protein of interest. PCFPs are a subfamily of fluorescent proteins that undergo an excitation and emission spectra shift upon stimulation with a specific wavelength of light (38). This approach enables the photoconversion of a small subpopulation of the PCFP-tagged protein, allowing for specific tracking of the photoconverted protein fluorescence intensity as a direct readout of both protein localization changes and degradation (after accounting for photobleaching effects). This approach also eliminates the need to average bulk cell responses as with cycloheximide chase assays, enabling investigation of how pHi regulates protein degradation in single cells.

To monitor live-cell β-catenin protein degradation rates, we selected mMaple3, a PCFP that transitions from green-to-red fluorescence upon exposure to 405 nm light (39). Derived from the parent protein mMaple, mMaple3 has reduced dimerization tendency, superior brightness in the converted channel, and improved photoconversion efficiency (defined as the proportion of protein that yields detectable fluorescence signal) (39–41). To validate that we can measure differences in protein degradation rates of photoconverted mMaple3 using confocal microscopy, we transiently expressed mMaple3 alone or N-terminally fused to β-catenin (mMaple3-β-catenin) or a degradation resistant β-catenin mutant (H36R-β-catenin) (5).

We hypothesized that this approach would allow for comparisons of relative protein degradation rates in single cells across treatment conditions. We first tested whether we could observe protein half-life differences in MDCK cells expressing mMaple3 alone, wild-type β-catenin, or a degradation-resistant β-catenin mutant (H36R-β-catenin). We photoconverted the target protein using a spatially restricted region of interest (Supplemental Video 1, β-catenin). We calculated normalized mean fluorescence intensity decay curves from cells and found that wild-type β-catenin was lost more rapidly than either H36R-β-catenin or mMaple3 alone (**Figure 5A**). It is important to note that this assay does not permit the determination of absolute protein half-life, as loss of fluorescent signal in this assay includes protein degradation, diffusion, and photobleaching of the PCFP. Indeed, we found that all proteins (including mMaple3) exhibited a two-phase fluorescence decay curve with a rapid decay followed by a slower loss of signal (**Figure 5A**). All subsequent analyses used two-phase decay curve fits and reported half-life values are for the slow rate, as the fast rates were not different across the proteins studied and likely reflect effects of diffusion.

**Figure 5:**
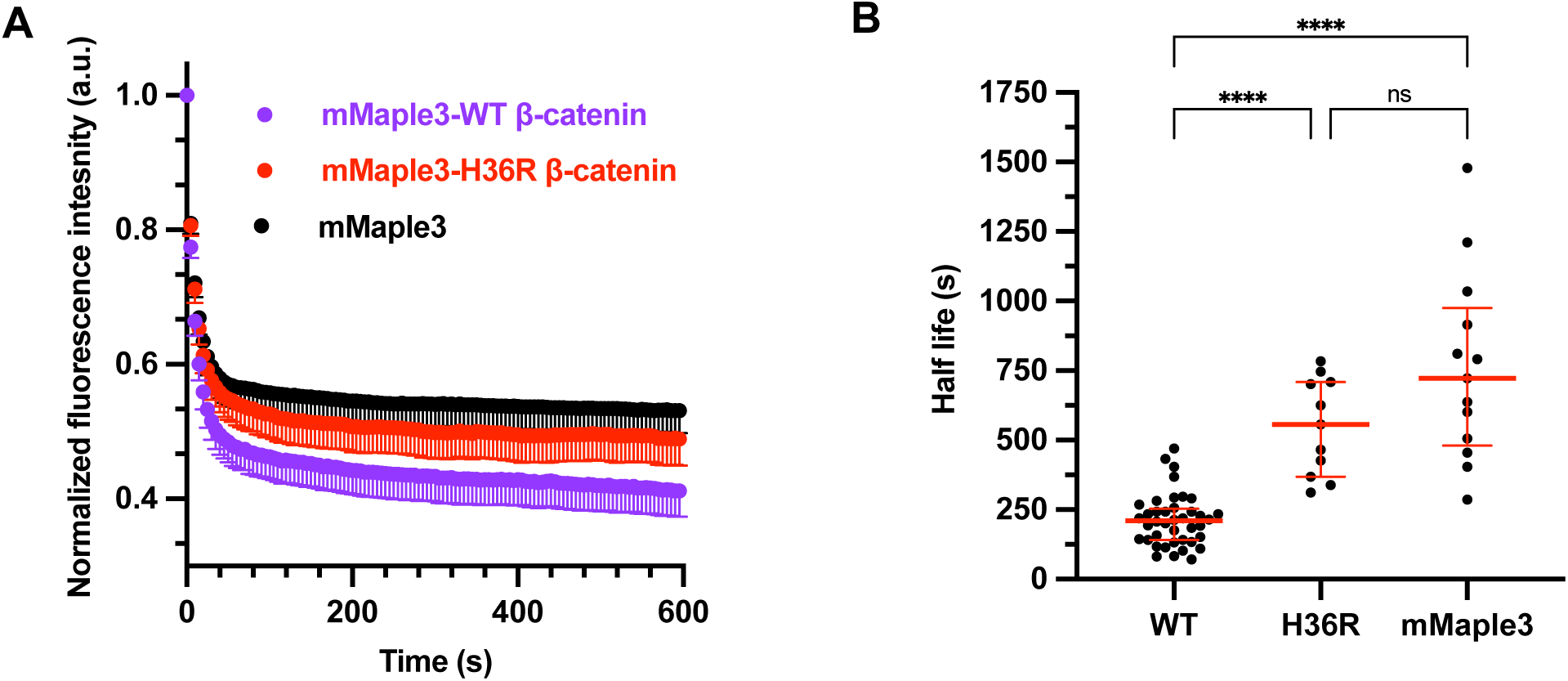
mMaple3 photoconversion can report on relative single-cell protein degradation rates. **(A)** Normalized fluorescence intensity after photoconversion of mMaple3-β-catenin, mMaple3-H36R-β-catenin, and mMaple3 alone. Shown are mean cell traces, error bars below mean indicate SEM. **(B)** Protein half-life was calculated or each cell in A and shown in a scatter plot (median±IQR) (mMaple3, n=12, N=3; mMaple3-H36R-β-catenin, n=11, N=3; mMaple3-β-catenin, n=19, N=4). Statistical significance in B was determined using Kruskal-Wallis test with Dunn’s correction for multiple comparisons. ****P<0.0001.

We next determined protein half-life from single-cell normalized fluorescence decay curves. Using this approach, we found that the median single-cell half-life of mMaple3 was 772 seconds, while the median single-cell half-life of mMaple3-β-catenin was significantly reduced to just 206 seconds (**Figure 5B**). This suggests that the shorter mMaple3-β-catenin half-life is being driven by ubiquitination and degradation of β-catenin in the fusion protein. Confirming this, a degradation-resistant mutant of β-catenin (mMaple3-H36R-β-catenin) had a significantly increased median half-life of 557 seconds in this assay, which was not statistically significantly different from that of mMaple3 alone (**Figure 5A, B**). Taken together, these data demonstrate mMaple3-β-catenin is i) degraded much faster than mMaple3 alone and ii) can be stabilized using a degradation-resistant mutant. Thus, these data validate the single-cell system as a reporter of mMaple3-β-catenin degradation dynamics in single cells and allow us to compare relative half-life values between pHi conditions. Next, we measured mMaple3-β-catenin degradation rates in single MDCK cells with pHi manipulation.

### Intracellular pH titrates the degradation rates of β-catenin in single epithelial cells

We next altered pHi in mMaple3-β-catenin expressing MDCK cells and photoconverted cytoplasmic mMaple3-β-catenin, enabling us to monitor relative pH-dependent degradation rates of cytoplasmic β-catenin in single cells. We observed a reduced mean half-life at high pHi (137.1s) and increased mean half-life at low pHi (265.3s) compared to control MDCK cells (209.9s) (**Figure 6A**). We also performed experiments using the stabilized β-catenin mutant (H36R-β-catenin) and found that the mean half-life of mMaple3-H36R-β-catenin was increased across all pHi conditions compared to mMaple3-β-catenin (**Figure 6B**). This suggests that turnover of WT β-catenin is faster than the stabilized mutant under all pHi conditions.

**Figure 6.**
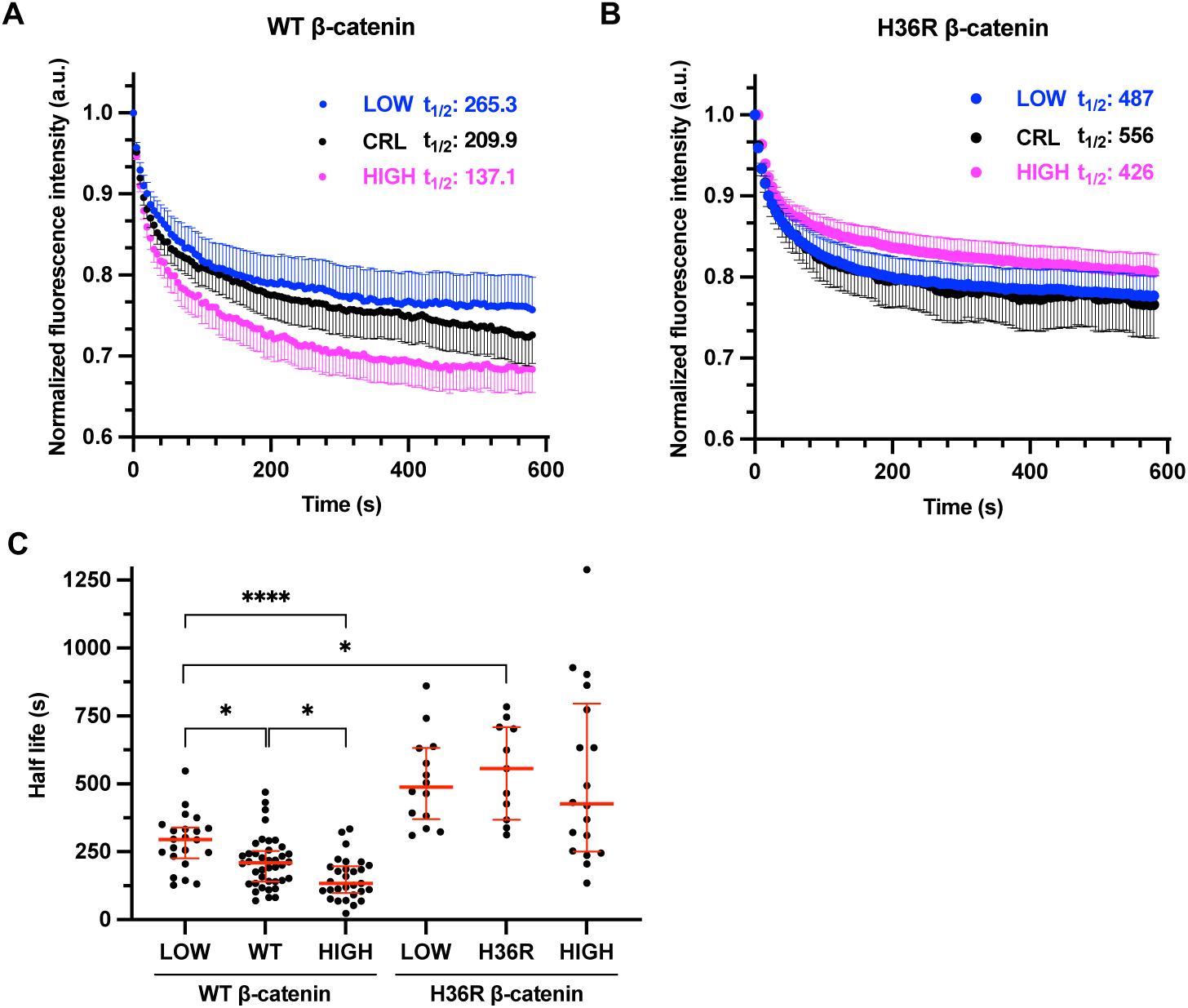
mMaple3-β-catenin has pH-dependent degradation, while mMaple3-H36R-β-catenin is stabilized and pH-insensitive. **(A)** Normalized fluorescence intensity traces from MDCK cells expressing mMaple3-β-catenin under low, control, and high pHi conditions. Dots show mean cell traces, error bars indicate SEM. Reported protein half-life calculated from mean cell traces. LOW, n=14; CRL, n=9; HIGH, n=20 from 3 biological replicates. **(B)** Normalized fluorescence intensity traces from MDCK cells expressing mMaple3-β-catenin under low, control, and high pHi conditions. Dots show mean cell traces, error bars below (CRL) or above (HIGH, LOW) mean indicate SEM. Reported protein half-life calculated from mean cell traces. LOW, n=14; CRL, n=11; HIGH, n=18 **(C)** Single-cell protein half-life values were obtained by fitting individual single-cell traces from cells collected as in A and B. (median±IQR) Outliers were removed using the ROUT method (Q=1%). WT LOW, n=22; WT CRL, n=40; WT HIGH, n=29 from 6 biological replicates. H36R LOW, n=14; H36R CRL, n=11; H36R HIGH, n=18 from 3 biological replicates. Statistical significance in C was determined using Kruskal-Wallis test with Dunn’s correction for multiple comparisons. *P<0.05, ****P<0.0001.

We next calculated single-cell half-life of mMaple3-β-catenin and found that low pHi significantly increased median mMaple3-β-catenin single-cell half-life compared to control and high pHi conditions while high pHi significantly reduced median mMaple3-β-catenin single-cell half-life compared to control cells and low pHi (**Figure 6C**). Importantly, median single-cell half-life of mMaple3-H36R-β-catenin was pHi-insensitive across all conditions tested (**Figure 6C**). This confirms prior work showing His36 is essential for pHi-dependent β-catenin stability (5). As a control, we also assessed mMaple3 stability in MDCK cells with and without pHi manipulation and found no difference in mean half-life or median single-cell half-life for mMaple3 (**Figure S8**). These experiments demonstrate that pHi titrates the degradation rate of cytoplasmic β-catenin in single cells, with faster degradation at high pHi and slower degradation at low pHi compared to control. These results also confirm that His36 is critical for pHi-dependent stability of β-catenin in single living cells.

In the photoconversion experiments described above, we performed cytoplasmic photoconversion and specific tracking of the cytoplasmic pool. This approach also allows us to monitor subcellular redistribution of photoconverted cytoplasmic mMaple3-β-catenin into the nucleus under altered pHi conditions. We measured nuclear accumulation of photoconverted mMaple3-β-catenin following cytoplasmic photoconversion and observed significantly increased nuclear accumulation of mMaple3-β-catenin at low pHi compared to high pHi cells (**Figure 7**). These data suggest that pHi modulates nuclear accumulation of photoconverted β-catenin and supports the observation that nuclear β-catenin abundance is increased at low pHi compared to control and high pHi (**Figure 2B**).

**Figure 7.**
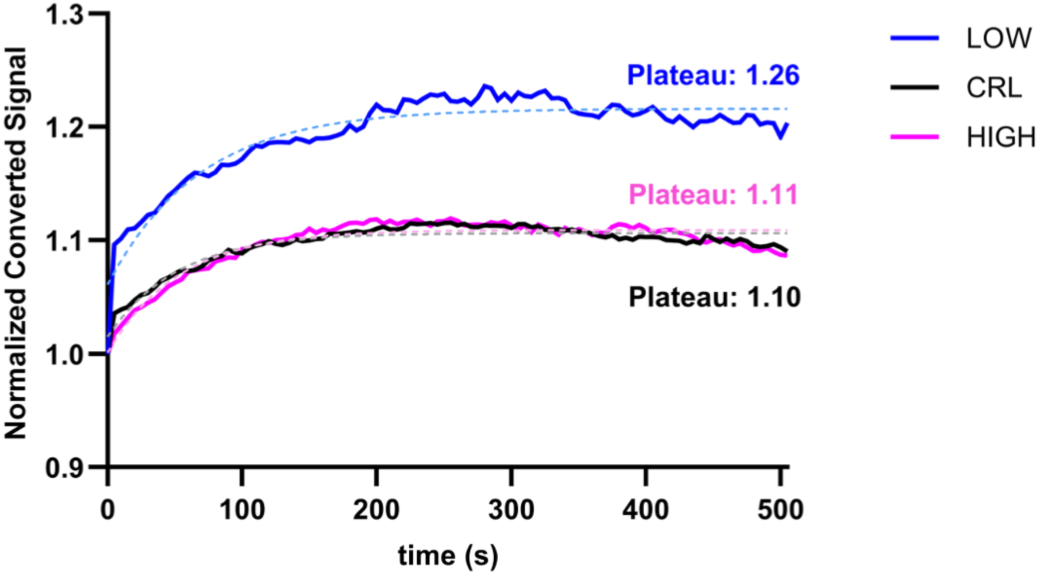
Low pHi increases nuclear accumulation of cytoplasmic photoconverted mMaple3-β-catenin. Normalized median cell traces for each condition are plotted and were fit using nonlinear regression (fitted curves shown as dashed, tinted lines) to obtain plateau values. LOW, n=14; CRL, n=13; HIGH, n=11; N=3. Statistical comparison was performed by determining whether a single curve could adequately fit both datasets, the null hypothesis was rejected (P<0.0001).

Our work suggests that low pHi significantly increases β-catenin abundance in the cytoplasm and nucleus due to attenuated cytoplasmic degradation. A remaining unanswered question is whether the transcriptional activity of wild-type, endogenous β-catenin is altered as a consequence of pH-dependent degradation of cytoplasmic protein pools, as this was not addressed in prior work investigating pH-dependent β-catenin stability (5). To address this, we tested the hypothesis that low pHi increases the transcriptional activity of endogenous β-catenin, whereas high pHi reduces transcriptional activity.

### Decreased pHi is sufficient to increase transcriptional activity of β-catenin

While the function of β-catenin is regulated by its localization, recent work revealed that nuclear localization of total protein alone is not sufficient to determine β-catenin function (42). Thus, to further support these data, we assessed nuclear abundance of non-phosphorylated (transcriptionally active) β-catenin in single cells with altered pHi. As expected, immunofluorescence staining for active β-catenin (**Figure 8A**) showed that active β-catenin enriched at cell membranes and is increased in the nucleus of MDCK cells at low pHi compared to control and high pHi cells (**Figure 8B**). These data demonstrate that low pHi increases transcriptionally-active forms of β-catenin in the nucleus, suggesting that pHi-dependent regulation of β-catenin abundance may also regulate transcriptional activity.

**Figure 8:**
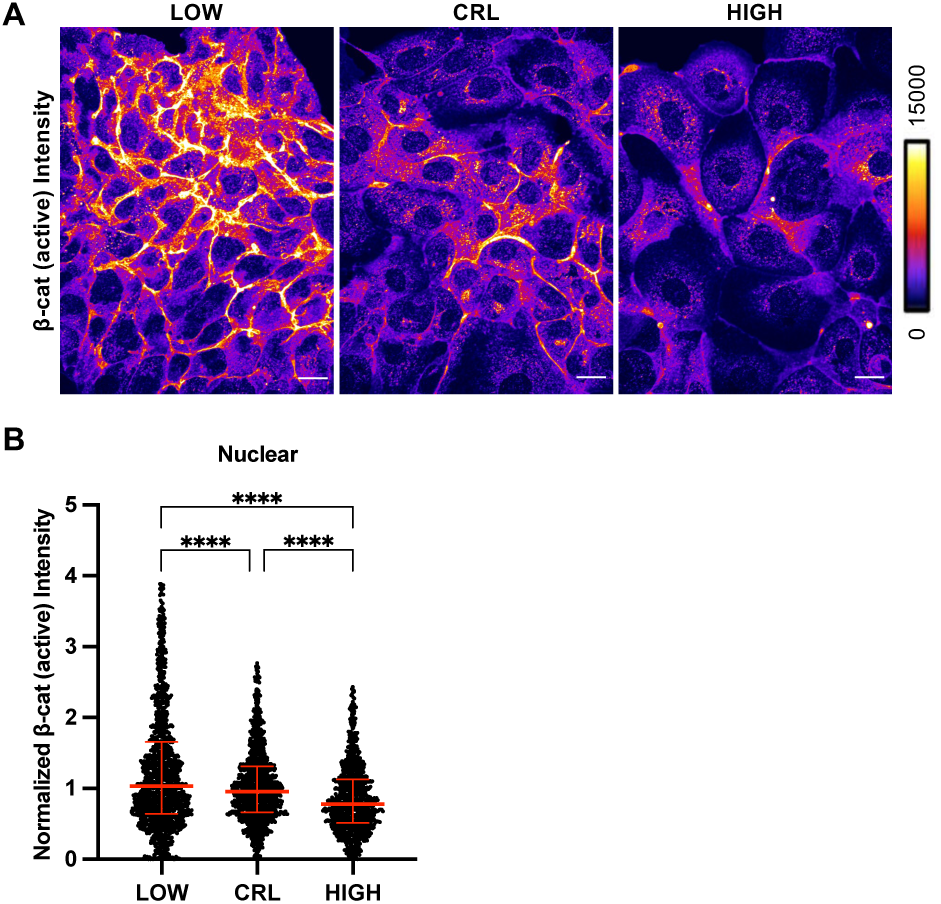
Active β-catenin is increased in the nucleus at low pHi compared to control and high pHi. **(A)** Representative confocal images of MDCK cells fixed and stained for non-phosphorylated (active) β-catenin (Ser33/Ser37/Thr41). β-catenin is pseudocolored according to scale. Scale bar: 20 µM. **(B)** Quantification of nuclear intensities, normalized to the median of control conditions within each biological replicate. Individual cells are shown as gray dots, (n=756-1231, from 4 biological replicates). Medians and interquartile ranges shown in red. Statistical significance for B and C as determined by Kruskal-Wallis test with Dunn’s multiple comparisons correction. ****P<0.0001.

To test this hypothesis, we next performed single-cell analysis of pH-dependent β-catenin transcriptional activity using a highly specific LEF-1/TCF transcriptional reporter plasmid that drives expression of destabilized GFP (TOPdGFP) (44). We generated a stable clonal MDCK cell line expressing this reporter (see methods). We first validated that the reporter cell line responded to Wnt3a stimulation across a range of Wnt3a concentrations (**Figure 9A, B**).

**Figure 9:**
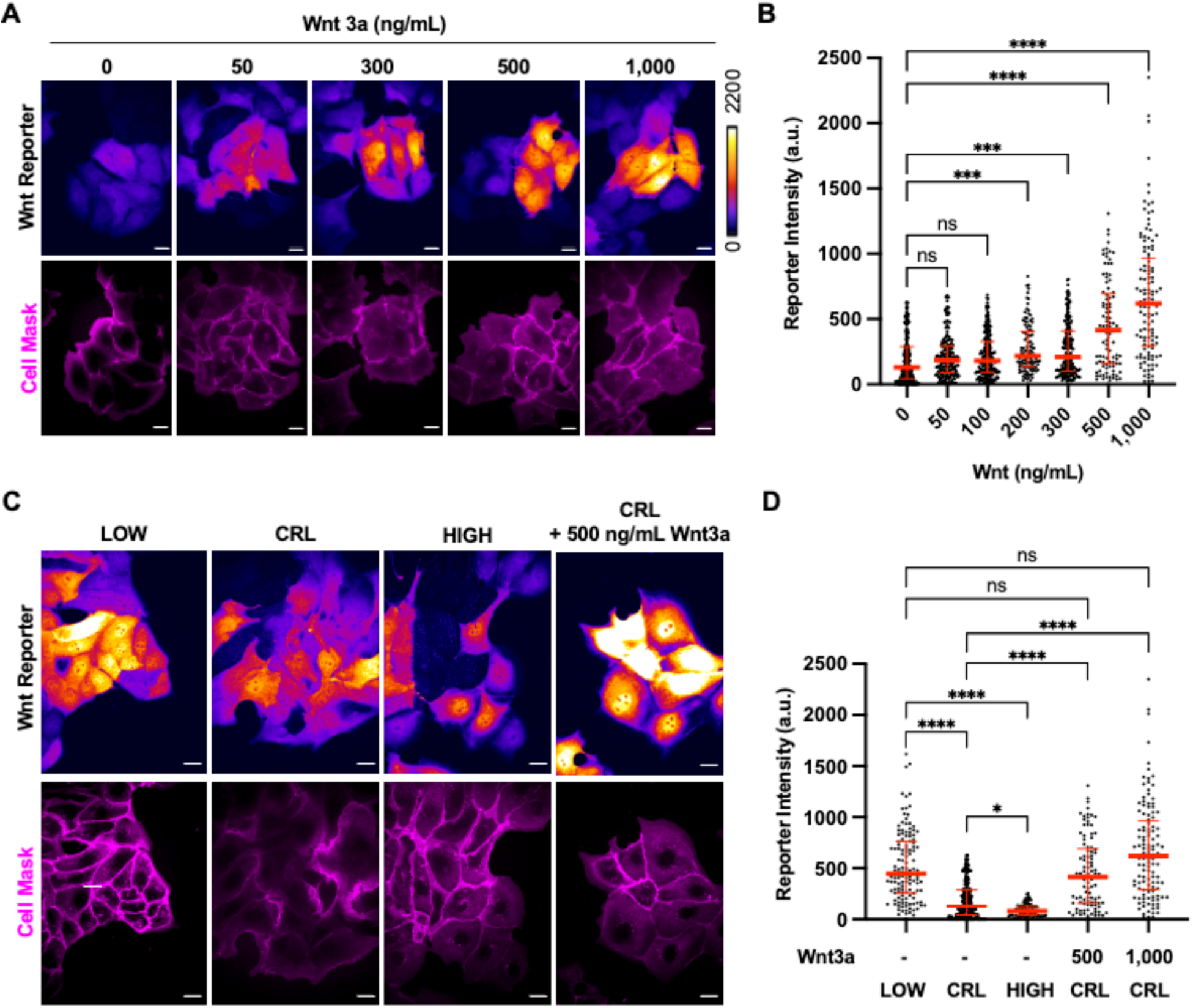
Low pHi phenocopies Wnt3a-mediated activation of β-catenin transcription. **(A)** Representative images of control MDCK cells expressing a TOPdGFP reporter with cell membranes visualized with Cell Mask (magenta). Cells were treated with Wnt3a at the indicated concentrations (0 to 1,000 ng/mL) for 24 hours. Scale bar = 20µm. **(B)** Quantification of single-cell reporter intensities from cells prepared as in A. Individual cells are shown as block dots with median and interquartile ranges shown in red lines; n=97-240 cells across 3 biological replicates. **(C)** Representative images of MDCK cells expressing a TOPdGFP reporter plasmid with and without pHi manipulation. Intensity shown according to scale, with cell membranes visualized with Cell Mask (magenta). **(D)** Quantification of single-cell reporter intensities from cells prepared as in C. Individual cells are shown as block dots with median and interquartile ranges shown in red lines; n=86-240 cells across 3 biological replicates. For B and D, statistical significance was determined using Kruskal-Wallis test with Dunn’s correction for multiple comparisons. *P<0.05, ***P<0.001, ****P<0.0001.

We then manipulated pHi for 24 hours prior to live-cell imaging of reporter fluorescence using confocal microscopy (**Figure 9C**). We observed a statistically significant increase in reporter fluorescence at low pHi compared to high pHi (**Figure 9D**). Importantly, lowering pHi produced reporter fluorescence that was statistically indistinguishable from the highest concentrations of Wnt3a tested (**Figure 9D**). We also found that high pHi reduced transcriptional activation of β-catenin compared to control (**Figure 9C,D**). This demonstrates that high pHi can reduce basal β-catenin transcriptional activity, even in the absence of Wnt-3a stimulation. These single-cell analyses show that low pHi alone can recapitulate the Wnt-active state in MDCK cells, validating low pHi as a sufficient driver of endogenous β-catenin transcriptional activity in normal epithelial cells.

## Discussion

Our data highlight an important role for pHi in serving as a rheostat to regulate β-catenin abundance, where low pHi increases β-catenin stability, which increases β-catenin co-localization to E-cadherin and drives increased transcriptional activity even in the absence of extracellular Wnt signal. Conversely, high pHi reduces β-catenin stability, which reduces β-catenin at E-cadherin-containing AJs and lowers β-catenin transcriptional activity compared to control cells. Our work significantly expands on prior work that only explored the effects of increased pHi on β-catenin abundance and did not fully characterize pH-dependent regulation of β-catenin function. Our work demonstrates that pHi changes within the physiological range can regulate the degradation, abundance, and function of endogenous β-catenin.

We also found that subcellular pools of β-catenin were differentially regulated by pHi. We show that pHi-mediated loss of β-catenin from E-cadherin-containing AJs can be rescued by plakoglobin to maintain AJ integrity at high pHi. While genetic knockdown of β-catenin was previously shown to recruit plakoglobin to adherens junctions (34), we demonstrate that loss of β-catenin at high pHi is sufficient to produce the same compensatory phenotype at AJs. We also show that at low pHi, β-catenin is increased and plakoglobin is reduced at E-cadherin containing AJs.

Taken together, these data suggest pHi dynamics can directly remodel AJ composition in normal epithelial cells. We have demonstrated that plakoglobin is able to rescue the loss of β-catenin at AJs at high pHi. Additionally, we have found that the composition of AJs changes with pHi, with a preference for β-catenin at low pHi and a preference for plakoglobin at high pHi. This presents a potential novel mechanism of pH-dependent AJ remodeling that may be a previously unrecognized reinforcer of pH-dependent cell polarity (45) or EMT (46). Moreover, the increased ratio of plakoglobin to E-cadherin at high pHi may also be indicative of downstream changes in desmosome formation at high pHi, as plakoglobin-containing AJs are a prerequisite for their establishment (47). Future work will explore the molecular mechanisms of pH-dependent AJ remodeling, characterize how the physical properties of AJ change with pHi, and expand on the functional output of this remodeling in the context of 3D tissue formation.

Collectively, our population and single-cell level analyses of β-catenin transcriptional activity demonstrate that modest stabilization of nuclear β-catenin is sufficient to drive significant changes in transcriptional activity. We found that low pHi increased β-catenin transcriptional activity and phenocopied transcriptional outcomes of Wnt3a treatment. Importantly, while plakoglobin can rescue the AJ function of β-catenin at high pHi, our data suggest plakoglobin cannot rescue the reduced signaling function of β-catenin at high pHi, as transcriptional activity at high pHi was significantly reduced compared to control. This aligns with prior work showing β-catenin/TCF4 complexes are transcriptionally active but that plakoglobin/TCF4 complexes are unable to bind DNA (48, 49). Our data show that pHi directly regulates β-catenin signaling and transcriptional activity in the absence of Wnt pathway activation and indicates that pH-dependent regulation of β-catenin signaling may indeed reinforce stem cell phenotypes previously associated with low pHi(4) and high Wnt signaling (50).

Others have shown that internalization of β-catenin: E-cadherin complexes upon growth factor stimulation causes increased transcriptional activity of β-catenin, suggesting β-catenin: E-cadherin complex formation is a prerequisite for signaling-competent β-catenin(51). Here, we observed that low pHi induced 1) an unexpected loss of junctional E-cadherin, 2) increased β-catenin in E-cadherin immune complexes from whole cell lysates and 3) increased transcriptional activity of β-catenin. Our data may suggest that at low pHi internalization of E-cadherin saturated with β-catenin may result in more transcriptionally active β-catenin re-localizing to the nucleus. While not directly tested in this work, synergy between enhanced E-β-catenin/E-cadherin internalization and slowed degradation of β-catenin may contribute to the robust signaling phenotype we observed. Future work from our lab will apply optogenetic tools to spatiotemporally manipulate pHi (52) while monitoring AJ protein dynamics.

Decades of research have been dedicated to understanding the dynamics and regulation of β-catenin in normal physiology and disease (8, 14). Our work provides new mechanistic insight in the context of pH-dependent differentiation, where stem cells have low pHi and high β-catenin abundance and pHi increases are necessary for differentiation, accompanied by the loss of β-catenin (4, 53). Future work will explore roles for pHi dynamics in regulating β-catenin abundance through pH-dependent stability to support both maintenance of pluripotency (stabilized β-catenin at low pHi) and facilitation of differentiation (destabilized β-catenin at high pHi).

Finally, our work may shed light on how pHi regulates β-catenin abundance and transcriptional activity in cancer. Constitutively increased pHi is an emerging hallmark of cancer that is associated with several cancer cell behaviors (25). While pHi has been shown to increase with metastatic potential in tissue-matched cell lines (3), spatial pHi heterogeneity exists in 3D cancer cell cultures (54, 55). Concurrent work in our lab has revealed that microenvironment can alter pHi and concomitantly modulate β-catenin abundance and acquisition of vasculogenic mimicry phenotypes in cancer cell models(56). Thus, our work suggests that pHi changes may serve as a key physiological cue to regulate β-catenin abundance and activity to drive transcriptional and cell adhesion changes associated with cancer cell adaptation. More work is needed to simultaneously quantify pHi and β-catenin activity in more complex 3D cancer models to determine if pH-dependent regulation of the Wnt pathway (through β-catenin regulation) alters tumorigenic phenotypes. Our work characterizes pHi dynamics as a key regulator β-catenin abundance, adherens junction composition, and signaling function and provides a foundation to better understand context-specific avenues for pHi-dependent β-catenin abundance in regulating normal and disease biology.

## Experimental procedures

### Cell Culture and Conditions

Madin-Darby Canine Kidney (MDCK) cells were cultured in EMEM (Quality Biological, 112-018-101) supplemented with 10% Fetal Bovine Serum (FBS; Peak Serum, cat: PS-FB3). Cells were maintained at 37⁰ C in a humidified incubator maintained at 5% CO_2_ (unless indicated otherwise). Cells were authenticated and tested for mycoplasma in November 2022 and again in December 2024.

### Intracellular pH manipulation

Cells were plated 24 hours prior to being treated for pHi manipulation. 24 hours after plating, the conditioned medium was aspirated and replaced with medium containing 5-(*N*-ethyl-*N*-isopropyl) amiloride) (Sigma, cat: A3085), an inhibitor of the sodium proton exchanger and S0859, a selective inhibitor of the sodium bicarbonate transporter (Sigma, cat: SML0638) 1 µM EIPA + 30 µM S0859 (Cf) (low pHi) or fresh medium and placed in an incubator maintained at 5% CO_2_ (low pHi and control) or 15% CO_2_ (high pHi) for 24 hours. Where indicated, proteasome activity was inhibited 18 hours after pHi manipulation treatment using 10 µM MG132 (Selleck, S2619) for a total of 6 hours prior to sample analysis or collection.

### Acquisition and analysis of microscopy images

Plates were imaged using a Nikon Ti-2 spinning disk confocal microscope with a stage-top incubator (Tokai Hit), a spinning disk confocal head (Yokogawa; CSU-X1), a Ti2-S-SE stage, multi-point perfect focus system (PFS), an Orca Flash 4.0 CMOS camera, four laser lines (405, 488, 561, 647 nm) with appropriate filter sets (DAPI: 405 nm laser excitation, 455/50 nm emission; GFP: 488 nm laser excitation, 525/36 nm emission; TxRed: 561 nm laser excitation, 605/52 nm emission; SNARF: 561 nm laser excitation, 705/72 nm emission; Cy5: 647 nm laser excitation, 705/72 nm emission) were used. Immunofluorescent and live-cell transcriptional reporter images were collected on a 40X oil immersion objective (Nikon, CFI PLAN FLUOR NA 1.30). Photoconversion images were collected on a 60X oil immersion objective (Nikon, CFI PLAN APO OIL 1.40). Live-cell imaging was performed after 24 hours of pHi manipulation. The stage top chamber was equilibrated to 37⁰ C and 5% CO_2_ (or 15% CO_2_ when capturing experimental images for high pHi) prior to loading plates on microscope.

### Trypan blue exclusion assay

MDCK cells were plated at 1×10^3^ cells per well in a 24-well plate. Cells were cultured for 24 hours after plating before being treated for pHi manipulation as described above. After the 24-hour treatment period, the medium was aspirated, the cells were washed with 1 mL pre-warmed 1X Dulbecco’s Phosphate-Buffered Saline (DPBS) (Quality Biological, cat: 114-057-101) prior to trypsinization. Cells were trypsinized in 100 µL of trypsin (Corning, cat: 25-053-CI) for 8-10 minutes to ensure detachment from the plates. Each well then received 100 µL pre-warmed culture medium to inactivate the trypsin and resuspend the trypsinized cells to a single-cell suspension. Immediately before counting cell suspensions, 50 µL of the suspension was mixed with 50 µL trypan blue 0.4% solution (Gibco, cat: 15250-061) in a separate Eppendorf tube. Cells were counted using 10 µL of the 1:1 cell suspension:trypan blue mixture in a hemocytometer for healthy (no dye uptake) and unhealthy/dying (blue) cells. Each biological replicate is the average of three wells (technical replicates) plated, treated, and counted on independent days.

### Population-level pHi measurement using BCECF-AM

MDCK cells were plated at 1×10^3^ cells/mL in 24-well plate 48 hours prior to performing the assay on the plate reader. Cells were treated for pHi manipulation as outlined above. After 24 hours of treatment, 2’-7’-Bis-(2-carboxyethyl)-5-(and-)-carboxyfluorescein, Acetoxymethyl Ester (BCECF-AM) (VWR, 891390244) was prepared as a 100 µM stock solution in 10% DMSO in DPBS. Cells were dye loaded with 1 µM BCECF-AM for 30 minutes in respective culturing conditions. Pre-warmed HEPES-based wash buffer (30 mM HEPES, 145 mM NaCl, 5 mM KCl, 10 mM glucose, 1 mM MgSO_4_, 1mM KHPO_4_, 2mM CaCl_2_, pH 7.4) and Nigericin buffer (25 mM HEPES, 105 mM KCl, 1 mM MgCl_2_ supplemented with 10 µM Nigericin (Fisher, N1495)) are prepared the day of the assay. Nigericin buffer is used to prepare three standard buffers with pH values ∼7.8, ∼7.2 and ∼6.5, where pH was accurately recorded to hundredths place for each point and each biological replicate for pHi back-calculation. After dye-loading, cells are washed 3×5 minutes with HEPES buffer containing the appropriate treatment at 37⁰ C. Experimental fluorescence intensities were measured on a Biotek Cytation5 plate reader (440ex/535em) and (490ex/535em) with kinetic reads taken every 39 seconds for 7 minutes (11 reads). HEPES wash buffer was aspirated and replaced with Nigericin buffer, pH ∼7.8 and incubated for 7 minutes at 37⁰ C before starting the kinetic read. This procedure was repeated for the other two Nigericin buffers. A data reduction step calculates the mean 490/440 intensity for experimental and standardized pH reads where slope and y-intercept are then calculated using Nigericin standard buffer pH values to the hundredth place. Population level pHi values are back-calculated using experimental ratios and Nigericin standard curve values.

### Whole-cell lysate preparation & protein quantification

After pHi manipulation (see above section) plates were removed from the incubator and placed on ice. The media was aspirated, and the plates were washed two times with ice-cold DPBS. Plates were wrapped with parafilm and stored at −80⁰ C if not prepared immediately. To washed or thawed plates on ice, 100 µL of lysis buffer (500 mM NaCl, 10 mM EDTA, 500mM HEPES-free acid, 10 mM EGTA, 10% Triton X-100, and protease inhibitor cocktail (Pierce™ Protease Inhibitor Tablets, A32965; 1 tablet/50 mL lysis buffer), pH 7.4) was added to each plate. Cells were removed using cell scrapers and transferred to pre-chilled microfuge tubes, briefly vortexed, then incubated on ice for 15 minutes. Lysates were centrifuged at 14,000 x g for 15 minutes at 4⁰ C. Clarified lysates were isolated to separate microfuge tubes on ice. The Bradford assay was used to quantify the total protein content of lysates (57) using Coomassie Protein reagent (Thermo Scientific, cat: 1856209). Lysates were stored at −80⁰ C if not immediately being used for SDS-PAGE sample preparation.

### Immunoprecipitation

After pHi manipulation (see above sections) plates were removed from the incubator and placed on ice. The media was aspirated, and the plates were washed two times with ice-cold DPBS. For each plate, 250 µL of immunoprecipitation lysis buffer (50 mM Tris, 150 mM NaCl, 1 mM NaF, 1% Triton X-100, and protease inhibitor cocktail (Pierce™ Protease Inhibitor Tablets, A32965; 1 tablet/50 mL lysis buffer) (pH 7.5) for 15 min on ice. Cells were scraped, and clarified lysate (10,000 rpm; 10 min) was used immediately. For E-cadherin immunoprecipitation, for each condition 100 µg of total protein was added to 2 µL of E-cadherin antibody (BD Bioscience, cat 610182) and incubated overnight at 4⁰ C with rotating. The next day, 40 µL of Protein G Dynabeads (Thermo Fisher, 10003D) was added to each sample and incubated for 2 hours at 4⁰ C with rotating. Beads were separated with a magnet and washed (3×5 minutes) with PBS, separating with magnet between washes. For elution, 2X Laemilli (non-reducing, BioRad 1610737) buffer was added to the bead volume, and beads were heated at 55⁰ C for 10 minutes before final magnetic separation. Eluted samples were then boiled for 10 minutes 100⁰ C with reducing agent (2-mercaptoethanol, 5% v/v) and then run on gels and transferred for immunoblotting (see Immunoblotting section for details).

### Immunoblotting

Protein samples were prepared in 6X Laemilli (reducing) buffer (Thermo Scientific, cat: J61337.AC) and boiled for 10 minutes at 100⁰ C. A total of 5 µg of MDCK lysate was added to each lane for immunoblot analysis. Protein samples were loaded onto 10% SDS-PAGE gels, which were run at 125V and 0.08 Amps for 15 minutes to stack samples. Protein samples were separated by running the gel at 175 V and 0.08 A for an hour and a half. Polyvinylidene difluoride (PVDF) membranes were pre-soaked in 100% ethanol for 5 minutes and rinsed 3X in water. Wet transfers were performed on ice at a constant amperage of 0.04 A (100V) for an hour and half in transfer buffer (Tris-Glycine buffer, 10% methanol, 0.01% SDS). Membranes were blocked for 1 hour in 5% milk in 1X Tris-buffered Saline + 0.1% Tween® 20 (TBS-T) at room temperature with shaking. For phospho-β-catenin blots, membranes were blocked in 3% BSA in TBS-T. Membranes were washed 3×5 min in 1X TBS-T before being cut and incubated with primary antibodies. The following primary antibodies were used: β-catenin (BD Biosciences, 610154; 1:5000 dilution in 3% BSA in 1X TBS-T), plakoglobin (Cell Signaling, 2309S; 1:2000 dilution in 3% BSA in 1X TBS-T), phospho β-catenin (pSer33/pSer37/pThr41) (Cell Signaling, 9561, 1:1,000 dilution in 1% BSA in 1X TBS-T), non-phospho(active)-β-catenin (Ser33/Ser37/Thr41) (Cell Signaling, 8814, 1:1,000 dilution in 1% BSA in 1X TBS-T), E-cadherin (Invitrogen, 13-1900, 1:1000 dilution in 3% BSA in 1X TBS-T), and actin (Santa Cruz, sc-58673; 1:500 diluted in 3% BSA in 1X TBS-T). Primary antibodies were incubated overnight at 4⁰ C with shaking. Primary antibody solutions were removed, and membranes were washed 3×5 minutes in 1X TBS-T before incubating secondary antibodies (Goat anti-mouse/rabbit HRP-conjugated antibodies; 1:10,000 diluted in 5% milk in 1X TBS-T) for 1 hour at room temperature with shaking. Secondary antibody solutions were removed, and membranes were washed 3×5 minutes in 1X TBS-T before being developed using Pierce SuperSignal West Pico PLUS (Thermo Scientific, cat: 34580) (β-catenin, actin, E-cadherin) or Pierce SuperSignal West Femto PLUS (Thermo Scientific, cat: 34095) (plakoglobin, phospho β-catenin, and all immunoprecipitation blots). Chemiluminescence and colorimetric images were acquired using a BioRad ChemiDoc imager.

### Western Blot Analysis

Raw files were exported from ChemiDoc imager and opened in ImageJ. Band intensities were acquired by densitometry analysis (area under the curve of individual bands in respective lanes). For each replicate blot, target protein intensities were normalized to the intensity of loading controls before normalizing each condition ratio to that of the control within biological replicates.

### Single-cell pHi calculations using SNARF-AM and confocal microscopy

Cells were plated at 5×10^3^ cells per well (4-well glass bottom dish, Matsunami, cat: D141400) or 2 x10^4^ cells per dish (35 mm dish with 14 mm glass cover slip, Matsunami, cat: D11130H) in complete medium 48 hours prior to imaging. Cells were treated for pHi manipulation as described above. Fields of view (FOVs) were selected by viewing cells in differential interference contract (DIC). Cells were dye loaded in conditioned media with 10 µM of 5-(and- 6)-Carboxy SNARF ™-1 Acetoxymethyl Ester, Acetate (Fisher Scientific, cat: C1272) prepared as a 100 µM stock solution dissolved in a 10% DMSO in DPBS for 10 minutes on the microscope stage. Dye-containing media was removed, and plates were washed three times with pre-warmed complete growth medium containing appropriate treatment. A 10-minute equilibration period was used prior to capturing images in SNARF (20% laser power, 400 ms), TxRed (20% laser power, 400 ms), and DIC (200 ms) channels for each FOV with five z-stacks (+/-1 µm; 0.5 µm steps). SNARF dye was calibrated using three pre-warmed Nigericin buffers (as described above). Complete medium was removed after the initial capture and the pH ∼7.8 Nigericin buffer was used to wash the plate and incubated for 10 minutes prior to capture. This procedure was repeated for standard points at pH ∼7.2 and ∼6.5. After data acquisition the NIS-Elements Analysis software was used to define single-cell regions of interest (ROIs) after background subtraction for each channel. The mean fluorescence intensity for each ROI in each channel was exported to Microsoft Excel. Ratios were calculated by dividing the SNARF mean intensity by the TxRed mean intensity for each capture. The three ratios for each nigericin point and their corresponding pH values were used to calculate slope and y-intercept. These values and the experimental ratio of SNARF/TxRed were used to back calculate single-cell pHi values.

### Immunofluorescent Staining

Cells were cultured in 35 mm imaging dishes containing a 14 mm glass coverslip and plated for experiments 48 hours prior to fixation. Cells were treated for pHi manipulation as described above. Intracellular pH was manipulated for 24 hours before removing culture dishes from the incubator and washing them with ice-cold DPBS. After being washed, cells were fixed in 3.7% paraformaldehyde in DPBS for 10 minutes at room temperature with shaking. The fixative solution was removed, and cells were washed 3×2 minutes in DPBS. Cells were permeabilized for 10 minutes at room temperature in a 0.1% Triton X-100 in 1X DBPS. Permeabilization buffer was removed and washed 3×2 minutes before blocking for 1 hour at room temperature in a 1% Bovine Serum Albumin (BSA) (Millipore Sigma, cat: 2910-OP) in DPBS solution. The blocking solution was removed and washed 3×2 minutes in DPBS. Primary antibodies (see Immunoblotting methods 1.3.7 for vendor information) were diluted in antibody solution (1% BSA, 0.1% Triton X-100 in DPBS) with the following dilutions: mouse anti-β-catenin (1:200), rabbit anti-plakoglobin (1:400), rabbit anti-non-phospho-β-catenin (1:100), and rat anti-E-cadherin (1:200). Primary antibody solutions were incubated in dishes at 4⁰ C overnight with rocking. After primary antibody incubation, the plates were washed 3×2 minutes prior to incubating Alexa-Fluor conjugated secondary antibodies diluted 1:1000 in antibody solution for 1 hour at room temperature while being protected from light. Secondary antibody solutions were removed and washed from the plates 3×2 minutes using DPBS. Cells were counterstained with Hoechst dye (DAPI; Thermo Scientific, cat: 62249) diluted 1:10,000 in antibody solution for 10 minutes at room temperature protected from light. Counterstain was removed and washed 3×2 minutes in DPBS. Imaging dishes were stored and imaged in DPBS. Image acquisition details are outlined above. Z-stacks were collected 2 µm above and below a relative z-plane that captured the center of the cell determined using the DAPI channel. Nuclear pools of proteins were identified using Nikon Elements Analysis software by auto-detecting individual regions of interest (ROI) in the DAPI channel that represent individual nuclei. Whole cell regions of interest were hand drawn using E-cadherin as a membrane marker to differentiate single-cells, drawing the ROI interior to E-cadherin staining to prevent inclusion of membrane signal. Nuclear ROIs were subtracted from hand-drawn whole cell ROIs to produce cytoplasmic ROIs, excluding both cell membranes and nuclei. The mean intensity for each channel was exported for each z-plane that had the largest mean intensity in the DAPI which indicated the center of the individual cells. Junctional protein levels were quantified using IMARIS software (Oxford Instruments, version 9.5.1) by generating surfaces based on E-cadherin intensity. Mean intensities for all channels within each surface were exported and analyzed for statistical significance using GraphPad Prism software.

### Quantifying Cell Area and Cell Volume

MDCK cells treated with pHi manipulation conditions were labeled with CellMask Deep Red (Thermo Fisher, C10046; 1:20,000) for 15 minutes. Cells were then imaged with Z-stacks collected encompassing the entire cell volume. For cell area calculation, a single z-plane from the center of the cell was analyzed in IMARIS. Individual cells were identified using the Cells module based on the Cell Mask signal (cell Membranes) and cell areas were exported. For cell volume calculations, a surface was generated in IMARIS using the CellMask signal and the complete Z-stack, filtration settings were adjusted such that the smallest cell diameter detected was 12.0 µm and cell membrane detail was 1.2 µm before cell volumes were exported. Area and volume data were analyzed for statistical significance using GraphPad Prism software.

### Cycloheximide chase assay

Cells were plated at 1.5×10^4^ cells per well in 6 well plate 48 hours prior to collecting lysates. 24 hours after plating, cells were treated for pHi manipulation as described above. Stock solutions of cycloheximide (VWR, 9427) were prepared fresh for each experiment. After 20 hours of pHi manipulation, all plates except time-0 samples were treated with 50 µg/mL cycloheximide and returned to the appropriate incubator. Immediately after treatment, lysates from time-0 plates were prepared as described previously and the remaining samples were lysed every 30 minutes for 2.5 hours. Immunoblotting and analysis of β-catenin and actin were performed as described above. The ratio of β-catenin to actin for each time point was normalized to time=0 h within each condition and for each replicate. Normalized data for each biological replicate was imported to Prism where one-phase decay was used to determine half-life values.

### Quantifying protein degradation and re-localization using mMaple3 constructs via photoconversion and confocal microscopy

MDCK cells were seeded at 3×10^6^ cells in 10 cm dishes and allowed to adhere to the dish for 8 hours. 8 hours after seeding, cells were transfected with 15 µg mMaple3 containing plasmid (mMaple3 alone, mMaple3-β-catenin WT, or mMaple3-β-catenin-H36R) using Lipofectamine 3000 according to the manufacturer’s instructions. Transfections proceeded for 18 hours prior to the removal of transfection media, rinsing of the plate with 10 mL of warm DPBS, and addition of 1 mL trypsin. Cells were trypsinized at 37⁰ C for 8-10 minutes. After all the cells were detached, complete medium was added to the dish to create a single cell suspension. Using the transfected single cell suspension, 35 mm imaging dishes were seeded at 2.5×10^4^ cell per dish and adhered for 8 hours. After the cells adhered for 8 hours, pHi manipulation was performed as described above. Images were collected as described above. Cell nuclei were visualized via SPY650-DNA (Cytoskeleton, cat. #: CY-SC501) probe that was incubated for 2 hours at 37⁰ C at a 1:1,000 dilution in complete media. Photoconversion was performed using a digital micromirror device (DMD) patterned illumination system (Polygon 4000) at 405 nm. For each experiment, an extra plate was prepared to calibrate the DMD device for a fixed stimulation ROI with a conserved area across cells and conditions (i.e. 6×16 pixels). Pre-stimulation images were acquired using GFP (pre-conversion mMaple3; 30% laser power, 500 ms) and Cy5 (DNA; 25% laser power, 200 ms), which were used to place the stimulation ROI. Within the Nikon Elements software, an ND Stimulation protocol was used to stimulate at 405 nm within a user-defined stimulation ROI with a conserved area across cells and conditions (i.e. 6×16 pixels) and then acquire images across the entire FOV using TxRed (converted mMaple3; 30% laser, 500 ms). For these experiments, a pre-conversion image was acquired followed by photoconversion in the stimulation ROI of mMaple3 or mMaple3-β-catenin constructs using 405 nm light at 100% LED power for 90 seconds. Then the converted (TxRed) signal was monitored every 30 seconds for 10 minutes post-conversion. ROIs for nuclei and cytoplasm were generated using GFP (unconverted mMaple3) and Cy5 (DNA probe) images and applied to the photoconverted movies to track nuclear and cytoplasmic signal over time.

Cytoplasmic tracking data were exported to excel and normalized to the first capture following photoconversion. Single-cell traces were then fit using nonlinear regression, comparing one-phase and two-phase decay curve fits with a constrained Y_o_ equal to 1.0. For all conditions and proteins, two-phase decay fit the data more accurately, but revealed similar values for the fast rate of decay for mMaple3 alone, mMaple3-β-catenin WT, and mMaple3-H36R-β-catenin, indicating this represents diffusion and/or bleaching. Single cell curves were then normalized to the time point after photoconversion that represented the start of the slow decay curves. This normalization accounts for the initial loss of fluorescent signal due to photobleaching and diffusion that was consistent across construct and conditions to enable curve fitting of protein stability. All reported half-life values are from the slow phase decay fits. The same two-phase decay models were used to compare mMaple3-β-catenin degradation under low, control, and high pHi conditions. Nuclear re-localization was quantified by generating nuclear ROIs based on SPY650-DNA intensity (see above) and quantifying nuclear intensity over the same time course used to monitor cytoplasmic degradation, in matched cells. Nuclear intensity was normalized to t=185 seconds and median traces were imported to Prism and fit using nonlinear regression and a one-phase association model with a constrained Y_o_ equal to 1.0 to account for the normalization we performed. Overall nuclear accumulation was readout by comparing plateau values between pHi conditions, where statistical comparison demonstrated that the same plateau value cannot adequately fit each dataset.

### Single-cell β-catenin transcriptional activity assays using live-cell microscopy

Lentiviral-TOP/FOP-dGFP-reporters (wild-type consensus plasmid TOP: Addgene plasmid #14715; http://n2t.net/addgene:14715; RRID:Addgene_14715; inverted consensus plasmid FOP: Addgene plasmid # 14885; http://n2t.net/addgene:14885 ; RRID:Addgene_14885) were a gift from Tannishtha Reya. For single-cell transcriptional assay, we conducted preliminary experiments using transient transfections. Cells were seeded at 3×10^6^ cells in 10 cm dishes and allowed to adhere to the dish for 8 hours. 8 hours after seeding, cells were transfected with 0 µg (mock transfection) or 15 µg TOP-dGFP-reporter/FOP-dGFP-reporter plasmid using Lipofectamine 3000 according to the manufacturer’s instructions. Transfections proceeded for 18 hours prior to the removal of transfection media, rinsing of the plate with 10 mL of warm DPBS, and addition of 1 mL trypsin. Cells were trypsinized at 37⁰ C for 8-10 minutes. After all the cells were detached, complete medium was added to the dish to create a single cell suspension. Using the transfected single cell suspension, 35 mm imaging dishes were seeded at 1.5×10^4^ cell per dish and adhered for 8 hours. After the cells adhered for 8 hours, the plating medium was removed and replaced with complete media used for pHi manipulations (see section above) or Wnt stimulation (40 ng/mL (Cf) murine Wnt3a (Peprotech; cat: 315-20)).

After validating the reporter, a stable cell line was generated. First, the TOP-dGFP reporter plasmid was transfected into MDCK cells. Briefly, single-cell dilutions of transfected MDCK cells were plated at low density (50 cells/mL) in 96-well plates. Colonies were tested for TOP-dGFP expression in a Wnt stimulation protocol (with and without 1000 ng/mL murine Wnt3a (Peprotech; 315-20)). A final clone was chosen that had robust reporter response and with matched morphology and pHi of parentals. Cells were seeded at 1.5×10^4^ cell per 35 mm imaging dish and adhered for 8 hours. After adherence, plating medium was removed and replaced with complete media for pHi manipulations (see section above) with or without Wnt stimulation (0 ng/mL to 1,000 ng/mL (Cf) murine Wnt3a (Peprotech; 315-20)).

For all single-cell Wnt stimulation assays, images were acquired as outlined in the above sections. Cell nuclei and cell membranes were visualized via Hoechst dye (DAPI; Thermo Scientific, cat: 62249; 1:10,000) and CellMask Deep Red (Thermo Fisher, C10046; 1:20,000), respectively, incubated for 15 minutes at 37⁰ C in complete media. Fields of view were selected by visualizing nuclei (DAPI) and images were collected in the DAPI (20% laser power, 400 ms), GFP (30% laser power, 500 ms), Cy5 (20% laser power, 400 ms), and DIC (32.6 DIA, 200 ms) channels. Whole-cell regions of interest (ROIs) were drawn within individual cells using cell mask as a membrane marker and the average GFP intensity for individual cells were exported to Excel. After stable cell line of TOP-dGFP, all cells were analyzed for GFP fluorescence intensity. For transient transfections, single-cell intensities that were greater than 2X the average intensity of mock transfected cells were imported to GraphPad Prism for statistical analysis and visualization. None of the cells expressing FOP-dGFP from the transient transfections met the 2-fold mock cutoff, indicating the robust specificity of this reporter assay.

### Statistics

GraphPad Prism (10.0) was used to prepare graphs and perform statistical comparisons as indicated in each figure legend. Normality tests were performed on all datasets after performing outlier analysis on pooled data (ROUT method; Q=1%), where normally distributed data is shown with means and non-normally distributed data is shown with medians. “N” indicates the number of biological replicates performed and “n” represents the number of technical replicates or individual measurements collected.

## Data Availability

The data associated with this work are included in this article and the supporting information file. Raw data are available upon request from the corresponding author.

## Supporting information

This article contains supporting information (Figures S1-S8).

## Acknowledgements

We would like to thank Tannishtha Reya for TOP/FOP-dGFP-reporter plasmids (TOP: Addgene plasmid #14715; http://n2t.net/addgene:14715; RRID:Addgene_14715; FOP: Addgene plasmid # 14885; http://n2t.net/addgene:14885; RRID:Addgene_14885). We would also like to thank members of the White lab for their helpful conversations and feedback during figure and manuscript preparation. The spinning disk confocal microscope used in this work was a part of the Notre Dame Integrated Imaging Facility (NDIIF) until Jan 2024.

## Author contributions

Conceptualization: BJC, KAW; Methodology: BJC, LNM, KAW; Validation: BJC, KAW; Formal analysis: BJC, LNM, KAW; Investigation: BJC, LNM, KAW; Resources: KAW; Data collection: BJC, LNM, KAW; Data curation: BJC, KAW; Writing – original draft: BJC, KAW; Writing - review & editing: BJC, LNM, KAW; Visualization: BJC, LNM, KAW; Supervision: BJC, KAW; Project administration: KAW; Funding acquisition: KAW

## References

1. Ulmschneider, B., Grillo-Hill, B. K., Benitez, M., Azimova, D. R., Barber, D. L., and Nystul, T. G. (2016) Increased intracellular pH is necessary for adult epithelial and embryonic stem cell differentiation. Journal of Cell Biology. 215, 345–355

2. Denker, S. P., and Barber, D. L. (2002) Cell migration requires both ion translocation and cytoskeletal anchoring by the Na-H exchanger NHE1. Journal of Cell Biology. 159, 1087– 1096

3. Spear, J. S., and White, K. A. (2023) Single-cell intracellular pH dynamics regulate the cell cycle by timing the G1 exit and G2 transition. Journal of Cell Science. 136, jcs260458

4. Liu, Y., Reyes, E., Castillo-Azofeifa, D., Klein, O. D., Nystul, T., and Barber, D. L. (2023) Intracellular pH dynamics regulates intestinal stem cell lineage specification. Nat Commun. 14, 3745

5. White, K. A., Grillo-Hill, B. K., Esquivel, M., Peralta, J., Bui, V. N., Chire, I., and Barber, D. L. (2018) β-Catenin is a pH sensor with decreased stability at higher intracellular pH. Journal of Cell Biology. 217, 3965–3976

6. Perez-Moreno, M., and Fuchs, E. (2006) Catenins: Keeping Cells from Getting Their Signals Crossed. Developmental Cell. 11, 601–612

7. Vleminckx, K., Kemler, R., and Hecht, A. (1999) The C-terminal transactivation domain of β-catenin is necessary and sufficient for signaling by the LEF-1/β-catenin complex in Xenopus laevis. Mechanisms of Development. 81, 65–74

8. Wodarz, A., and Nusse, R. (1998) Mechanisms of Wnt Signaling in Development. Annual Review of Cell and Developmental Biology. 14, 59–88

9. Chen, Y.-T., Stewart, D. B., and Nelson, W. J. (1999) Coupling Assembly of the E-Cadherin/β-Catenin Complex to Efficient Endoplasmic Reticulum Exit and Basal-lateral Membrane Targeting of E-Cadherin in Polarized MDCK Cells. Journal of Cell Biology. 144, 687–699

10. Huber, A. H., Stewart, D. B., Laurents, D. V., Nelson, W. J., and Weis, W. I. (2001) The Cadherin Cytoplasmic Domain Is Unstructured in the Absence of β-Catenin: A POSSIBLE MECHANISM FOR REGULATING CADHERIN TURNOVER *. Journal of Biological Chemistry. 276, 12301–12309

11. Drees, F., Pokutta, S., Yamada, S., Nelson, W. J., and Weis, W. I. (2005) α-Catenin Is a Molecular Switch that Binds E-Cadherin-β-Catenin and Regulates Actin-Filament Assembly. Cell. 123, 903–915

12. Yost, C., Torres, M., Miller, J. R., Huang, E., Kimelman, D., and Moon, R. T. (1996) The axis-inducing activity, stability, and subcellular distribution of beta-catenin is regulated in Xenopus embryos by glycogen synthase kinase 3. Genes Dev. 10, 1443–1454

13. Hart, M., Concordet, J.-P., Lassot, I., Albert, I., Santos, R. del los, Durand, H., Perret, C., Rubinfeld, B., Margottin, F., Benarous, R., and Polakis, P. (1999) The F-box protein β-TrCP associates with phosphorylated β-catenin and regulates its activity in the cell. Current Biology. 9, 207–211

14. Nusse, R., and Clevers, H. (2017) Wnt/β-Catenin Signaling, Disease, and Emerging Therapeutic Modalities. Cell. 169, 985–999

15. MacDonald, B. T., Tamai, K., and He, X. (2009) Wnt/β-Catenin Signaling: Components, Mechanisms, and Diseases. Developmental Cell. 17, 9–26

16. Mukherjee, A., Dhar, N., Stathos, M., Schaffer, D. V., and Kane, R. S. (2018) Understanding How Wnt Influences Destruction Complex Activity and β-Catenin Dynamics. iScience. 6, 13–21

17. Tetsu, O., and McCormick, F. (1999) β-Catenin regulates expression of cyclin D1 in colon carcinoma cells. Nature. 398, 422–426

18. He, T.-C., Sparks, A. B., Rago, C., Hermeking, H., Zawel, L., da Costa, L. T., Morin, P. J., Vogelstein, B., and Kinzler, K. W. (1998) Identification of c-MYC as a Target of the APC Pathway. Science. 281, 1509–1512

19. Moon, J. H., Lee, S. H., and Lim, Y. C. (2021) Wnt/β-catenin/Slug pathway contributes to tumor invasion and lymph node metastasis in head and neck squamous cell carcinoma. Clin Exp Metastasis. 38, 163–174

20. Daniels, D. L., and Weis, W. I. (2005) β-catenin directly displaces Groucho/TLE repressors from Tcf/Lef in Wnt-mediated transcription activation | Nature Structural & Molecular Biology. Nature Structural & Molecular Biology. 12, 364–371

21. Shahi, P., Seethammagari, M. R., Valdez, J. M., Xin, L., and Spencer, D. M. (2011) Wnt and Notch Pathways Have Interrelated Opposing Roles on Prostate Progenitor Cell Proliferation and Differentiation. Stem Cells. 29, 678–688

22. Scoville, D. H., Sato, T., He, X. C., and Li, L. (2008) Current View: Intestinal Stem Cells and Signaling. Gastroenterology. 134, 849–864

23. Benitez, M., Tatapudy, S., Liu, Y., Barber, D. L., and Nystul, T. G. (2019) Drosophila anion exchanger 2 is required for proper ovary development and oogenesis. Developmental Biology. 452, 127–133

24. Liu, Y., White, K. A., and Barber, D. L. (2020) Intracellular pH Regulates Cancer and Stem Cell Behaviors: A Protein Dynamics Perspective. Frontiers in Oncology. [online] https://www.frontiersin.org/articles/10.3389/fonc.2020.01401 (Accessed January 4, 2024)

25. White, K. A., Grillo-Hill, B. K., and Barber, D. L. (2017) Cancer cell behaviors mediated by dysregulated pH dynamics at a glance. Journal of Cell Science. 130, 663–669

26. Larsen, A. M., Krogsgaard-Larsen, N., Lauritzen, G., Olesen, C. W., Honoré Hansen, S., Boedtkjer, E., Pedersen, S. F., and Bunch, L. (2012) Gram-Scale Solution-Phase Synthesis of Selective Sodium Bicarbonate Co-transport Inhibitor S0859: in vitro Efficacy Studies in Breast Cancer Cells. ChemMedChem. 7, 1808–1814

27. Lee, D. H., and Goldberg, A. L. (1996) Selective Inhibitors of the Proteasome-dependent and Vacuolar Pathways of Protein Degradation in Saccharomyces cerevisiae. Journal of Biological Chemistry. 271, 27280–27284

28. Krieghoff, E., Behrens, J., and Mayr, B. (2006) Nucleo-cytoplasmic distribution of β-catenin is regulated by retention. Journal of Cell Science. 119, 1453–1463

29. Morgan, R. G., Ridsdale, J., Payne, M., Heesom, K. J., Wilson, M. C., Davidson, A., Greenhough, A., Davies, S., Williams, A. C., Blair, A., Waterman, M. L., Tonks, A., and Darley, R. L. (2019) LEF-1 drives aberrant β-catenin nuclear localization in myeloid leukemia cells. Haematologica. 104, 1365–1377

30. Valenta, T., Hausmann, G., and Basler, K. (2012) The many faces and functions of β-catenin. EMBO J. 31, 2714–2736

31. Grillo-Hill, B. K., Webb, B. A., and Barber, D. L. (2014) Chapter 23 - Ratiometric Imaging of pH Probes. in Methods in Cell Biology (Waters, J. C., and Wittman, T. eds), pp. 429–448, Quantitative Imaging in Cell Biology, Academic Press, 123, 429–448

32. Luciano, M., Versaevel, M., Vercruysse, E., Procès, A., Kalukula, Y., Remson, A., Deridoux, A., and Gabriele, S. (2022) Appreciating the role of cell shape changes in the mechanobiology of epithelial tissues. Biophysics Reviews. 3, 011305

33. Cowin, P., Kapprell, H.-P., Franke, W. W., Tamkun, J., and Hynes, R. O. (1986) Plakoglobin: A protein common to different kinds of intercellular adhering junctions. Cell. 46, 1063–1073

34. Fukunaga, Y., Liu, H., Shimizu, M., Komiya, S., Kawasuji, M., and Nagafuchi, A. (2005) Defining the roles of β-catenin and plakoglobin in cell-cell adhesion: Isolation of β-catenin/plakoglobin-deficient F9 cells. Cell Structure and Function. 30, 25–34

35. Wickline, E. D., Awuah, P. K., Behari, J., Ross, M., Stolz, D. B., and Monga, S. P. S. (2011) Hepatocyte γ-catenin compensates for conditionally deleted β-catenin at adherens junctions. Journal of Hepatology. 55, 1256–1262

36. Zhou, J., Qu, J., Xian, P. Y., Graber, K., Huber, L., Wang, X., Gerdes, A. M., and Li, F. (2007) Upregulation of γ-catenin compensates for the loss of β-catenin in adult cardiomyocytes. American Journal of Physiology - Heart and Circulatory Physiology. 292, 270–276

37. Kao, S.-H., Wang, W.-L., Chen, C.-Y., Chang, Y.-L., Wu, Y.-Y., Wang, Y.-T., Wang, S.-P., Nesvizhskii, A. I., Chen, Y.-J., Hong, T.-M., and Yang, P.-C. (2015) Analysis of Protein Stability by the Cycloheximide Chase Assay. Bio Protoc. 5, e1374

38. Shcherbakova, D. M., and Verkhusha, V. V. (2014) Chromophore chemistry of fluorescent proteins controlled by light. Current Opinion in Chemical Biology. 20, 60–68

39. Wang, S., Moffitt, J. R., Dempsey, G. T., Xie, X. S., and Zhuang, X. (2014) Characterization and development of photoactivatable fluorescent proteins for single-molecule-based superresolution imaging. Proceedings of the National Academy of Sciences of the United States of America. 111, 8452–8457

40. McEvoy, A. L., Hoi, H., Bates, M., Platonova, E., Cranfill, P. J., Baird, M. A., Davidson, M. W., Ewers, H., Liphardt, J., and Campbell, R. E. (2012) mMaple: A Photoconvertible Fluorescent Protein for Use in Multiple Imaging Modalities. PLOS ONE. 7, e51314

41. Zhang, L., Gurskaya, N. G., Merzlyak, E. M., Staroverov, D. B., Mudrik, N. N., Samarkina, O. N., Vinokurov, L. M., Lukyanov, S., and Lukyanov, K. A. (2007) Method for real-time monitoring of protein degradation at the single cell level. BioTechniques. 42, 446–450

42. Van Der Wal, T., and Van Amerongen, R. (2020) Walking the tight wire between cell adhesion and WNT signalling: A balancing act for β-catenin: A balancing act for CTNNB1. Open Biology. 10.1098/rsob.200267

43. Martínez-Estrada, O. M., Cullerés, A., Soriano, F. X., Peinado, H., Bolós, V., Martínez, F. O., Reina, M., Cano, A., Fabre, M., and Vilaró, S. (2006) The transcription factors Slug and Snail act as repressors of Claudin-1 expression in epithelial cells1. Biochemical Journal. 394, 449–457

44. Reya, T., Duncan, A. W., Ailles, L., Domen, J., Scherer, D. C., Willert, K., Hintz, L., Nusse, R., and Weissman, I. L. (2003) A role for Wnt signalling in self-renewal of haematopoietic stem cells. Nature. 423, 409–414

45. Frantz, C., Karydis, A., Nalbant, P., Hahn, K. M., and Barber, D. L. (2007) Positive feedback between Cdc42 activity and H+ efflux by the Na-H exchanger NHE1 for polarity of migrating cells. Journal of Cell Biology. 179, 403–410

46. Amith, S. R., Wilkinson, J. M., and Fliegel, L. (2016) Na+/H+ exchanger NHE1 regulation modulates metastatic potential and epithelial-mesenchymal transition of triple-negative breast cancer cells. Oncotarget. 7, 21091–21113

47. Lewis, J. E., Wahl, J. K., Sass, K. M., Jensen, P. J., Johnson, K. R., and Wheelock, M. J. (1997) Cross-Talk between Adherens Junctions and Desmosomes Depends on Plakoglobin. J Cell Biol. 136, 919–934

48. Roura, S., Miravet, S., Piedra, J., Herreros, A. G. de, and Duñach, M. (1999) Regulation of E-cadherin/Catenin Association by Tyrosine Phosphorylation *. Journal of Biological Chemistry. 274, 36734–36740

49. Simcha, I., Shtutman, M., Salomon, D., Zhurinsky, J., Sadot, E., Geiger, B., and Ben-Ze’ev, A. (1998) Differential nuclear translocation and transactivation potential of β- Catenin and plakoglobin. Journal of Cell Biology. 141, 1433–1448

50. Nusse, R. (2008) Wnt signaling and stem cell control. Cell Res. 18, 523–527

51. Howard, S., Deroo, T., Fujita, Y., and Itasaki, N. (2011) A Positive Role of Cadherin in Wnt/β-Catenin Signalling during Epithelial-Mesenchymal Transition. PLOS ONE. 6, e23899

52. Donahue, C. E. T., Siroky, M. D., and White, K. A. (2021) An Optogenetic Tool to Raise Intracellular pH in Single Cells and Drive Localized Membrane Dynamics. J. Am. Chem. Soc. 143, 18877–18887

53. Fevr, T., Robine, S., Louvard, D., and Huelsken, J. (2007) Wnt/β-Catenin Is Essential for Intestinal Homeostasis and Maintenance of Intestinal Stem Cells. Mol Cell Biol. 27, 7551– 7559

54. Dovmark, T. H., Saccomano, M., Hulikova, A., Alves, F., and Swietach, P. (2017) Connexin-43 channels are a pathway for discharging lactate from glycolytic pancreatic ductal adenocarcinoma cells. Oncogene. 36, 4538–4550

55. Tafech, A., Jacquet, P., Beaujean, C., Fertin, A., Usson, Y., and Stéphanou, A. (2023) Characterization of the Intracellular Acidity Regulation of Brain Tumor Cells and Consequences for Therapeutic Optimization of Temozolomide. Biology. 12, 1221

56. Lund, L. M., Marchi, A. N., Alderfer, L., Hall, E., Hammer, J., Trull, K. J., Hanjaya-Putra, D., and White, K. A. (2024) Intracellular pH dynamics respond to microenvironment stiffening and mediate vasculogenic mimicry through β-catenin. 10.1101/2024.06.04.597454

57. Bradford, M. M. (1976) A rapid and sensitive method for the quantitation of microgram quantities of protein utilizing the principle of protein-dye binding. Analytical Biochemistry. 72, 248–254

