## Supplemental Figures 1-8 for "Intracellular pH regulates β-catenin with low pHi increasing adhesion and signaling functions"

Supplementary Video 1. **Representative cytoplasmic photoconversion experiment of MDCK cells expressing mMaple3-β-catenin.** Scale bar = 20 µm. Video shown in inverse mono with stimulation ROI (red rectangle) overlaid each frame. The first frame represents pre-photoconversion image in channel to visualize converted mMaple3. The second frame shows converted mMaple3-β-catenin after 90 seconds of stimulation with 405 nm light within stimulation ROI with subsequent images collected over a 10-minute tracking period.

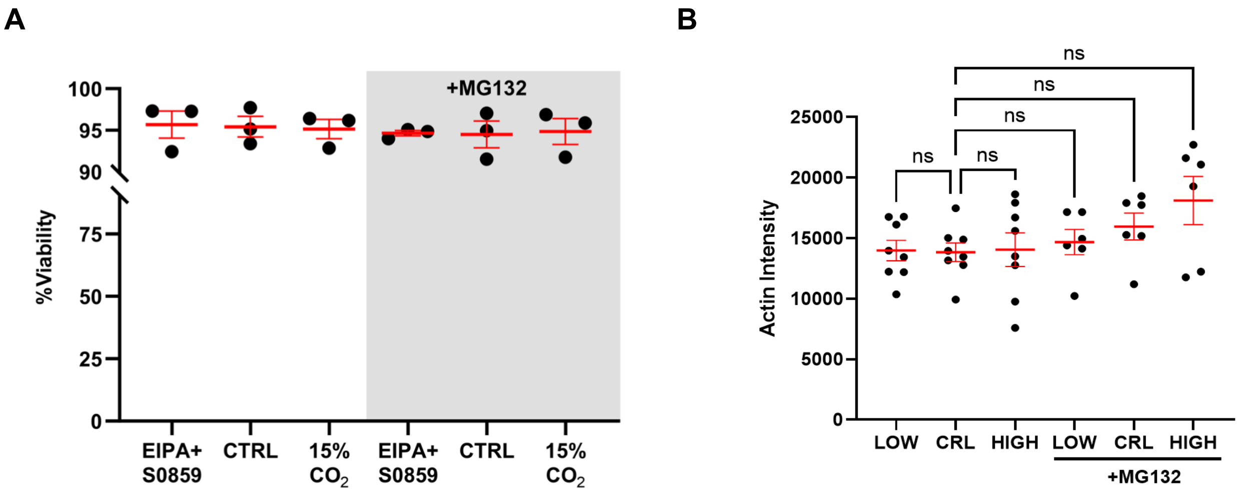


**Figure S1. Methods for manipulating intracellular pHi do not affect cellular viability or protein abundance of loading control proteins (Actin). (A)** Cellular viability as determined by trypan blue exclusion assay (see Methods). Means from 3 biological replicates were calculated from technical triplicates. Mean and SEM are shown. **(B)** Quantification of actin intensity from immunoblots prepared as in Figure 1. Mean and SEM are shown from 6-9 biological replicates. For both A and B, statistical significance was determined using one-way ANOVA test with Sidak's correction for multiple comparisons.

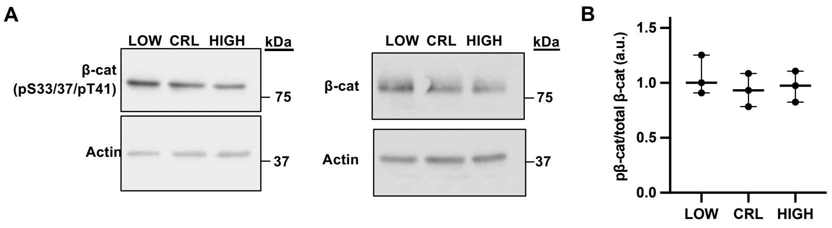


**Figure S2. Levels of phosphorylated β-catenin are not significantly altered by pHi in whole cell lysates. (A)** Representative immunoblot of β-catenin (β-cat), phospho-β-catenin (pSer31/pSer33/pThr41) and actin under low, control, and high pHi conditions in MDCK cells. **(B)** Quantification of β-catenin immunoblot data collected as described in A. Individual biological replicates were normalized to control MDCK within each experiment. Scatter plots show all data points from n=4 biological replicates. Statistical significance was determined using a ratio paired t-test between treatment groups and one-sample t-tests with a hypothetical mean of 1.0 when comparing to control (which had no variation due to normalization).

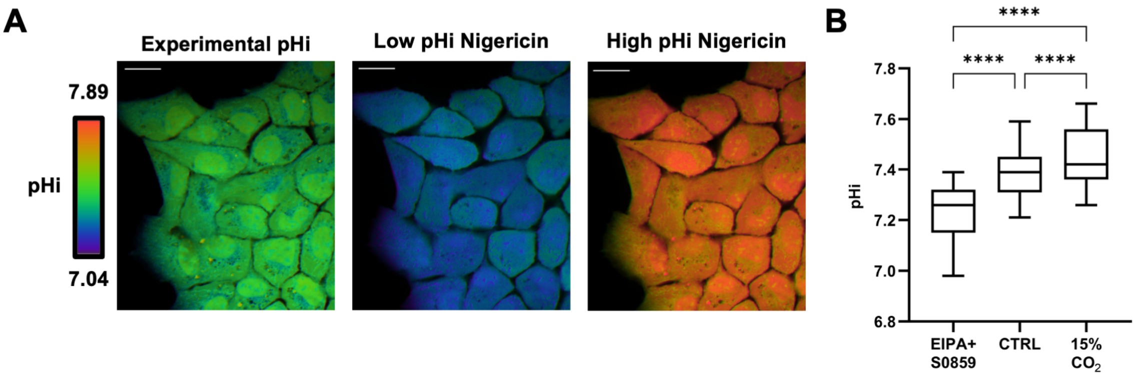


**Figure S3. 24-hour pHi manipulation conditions significantly alter pHi at the single-cell level. (A)** Representative confocal microscopy images of MDCK cells for pHi measurements and nigericin standardization (see Methods). Ratiometric display of emission at 640nm/580nm with 488 nm excitation. Scale bars: 20 μm. **(B)** Box and whisker plots of back calculated pHi (EIPA+S0859, n=244; CTRL, n=299; 15% $CO_2$, n=204; from two biological replicates). Medians are shown using Tukey display. Significance was determined by the Kruskal-Wallis test with Dunn's multiple comparisons correction. ****$P<0.0001$.

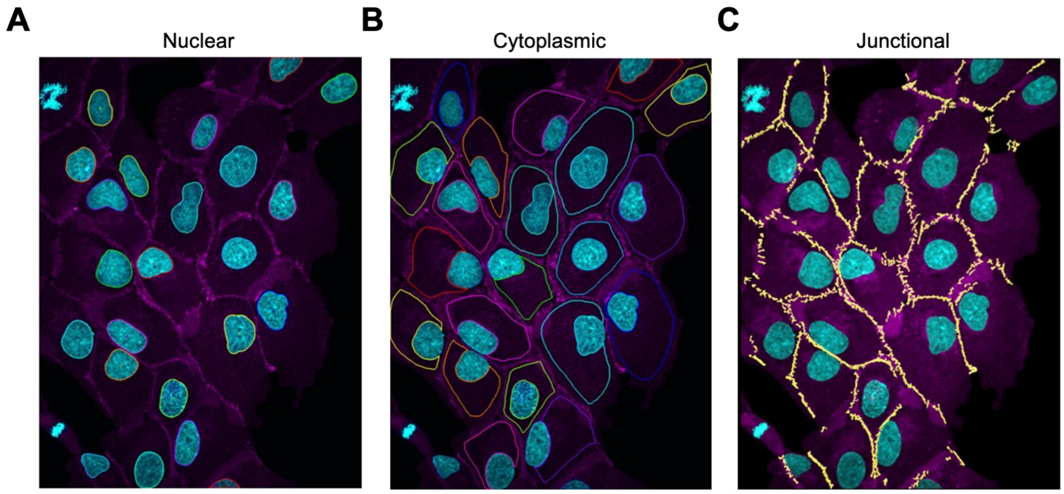


**Figure S4. Analysis pipeline for quantifying protein abundance in various subcellular pools. (A)** Nuclear regions of interest (ROIs) were generated by autodetecting single-cell nuclei based on Hoechst staining. **(B)** Cytoplasmic ROIs were created by subtracting nuclear ROIs (A) from whole cell ROIs by drawing interior to cell boundaries marked by E-cadherin staining (magenta). **(C)** 3D junctional surfaces were generated from z-stacks and thresholded based on E-cadherin (magenta) staining.

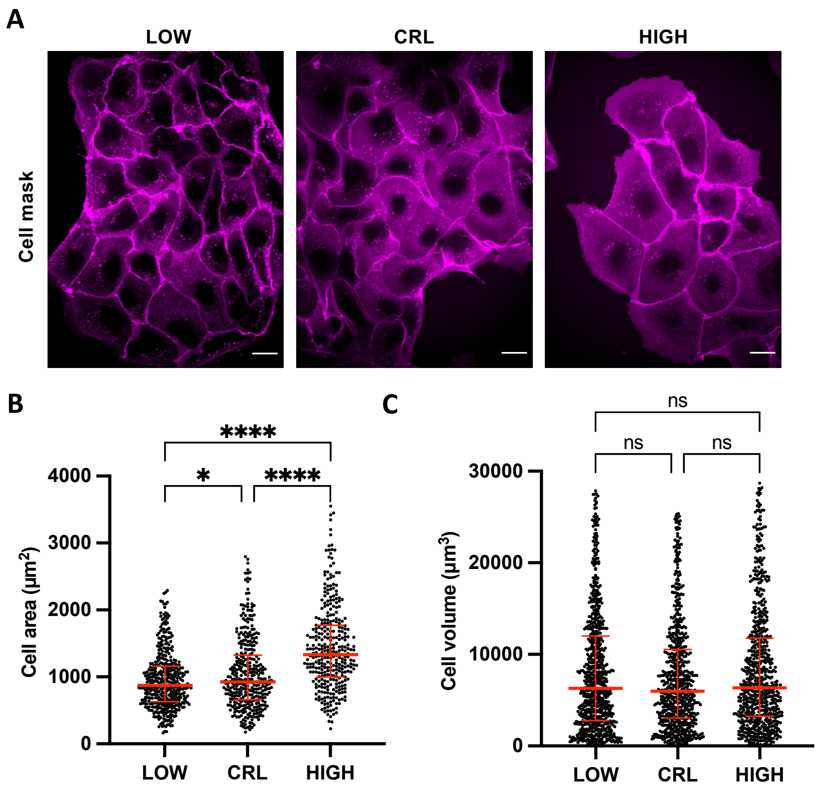


**Figure S5. Cell morphology analysis shows pH-dependent cell area changes but unchanged cell volume. (A)** Representative images of MDCK cells with low, control, and high pHi with membranes stained with CellMask Deep Red (see methods). Scale bars, 20 µm. **(B)** Quantification of single cell area for MDCK cells prepared as in A. Scatter plots of individual cells, median and interquartile range shown in red. N=3 biological replicates. **(C)** Quantification of single cell volume for MDCK cells prepared as in A. Scatter plots show individual cells, median and interquartile range shown in red. N=3 biological replicates. For B and C, significance was determined by the Kruskal-Wallis test with Dunn's multiple comparisons correction. $^*P<0.05$, $^{***}P<0.0001$.

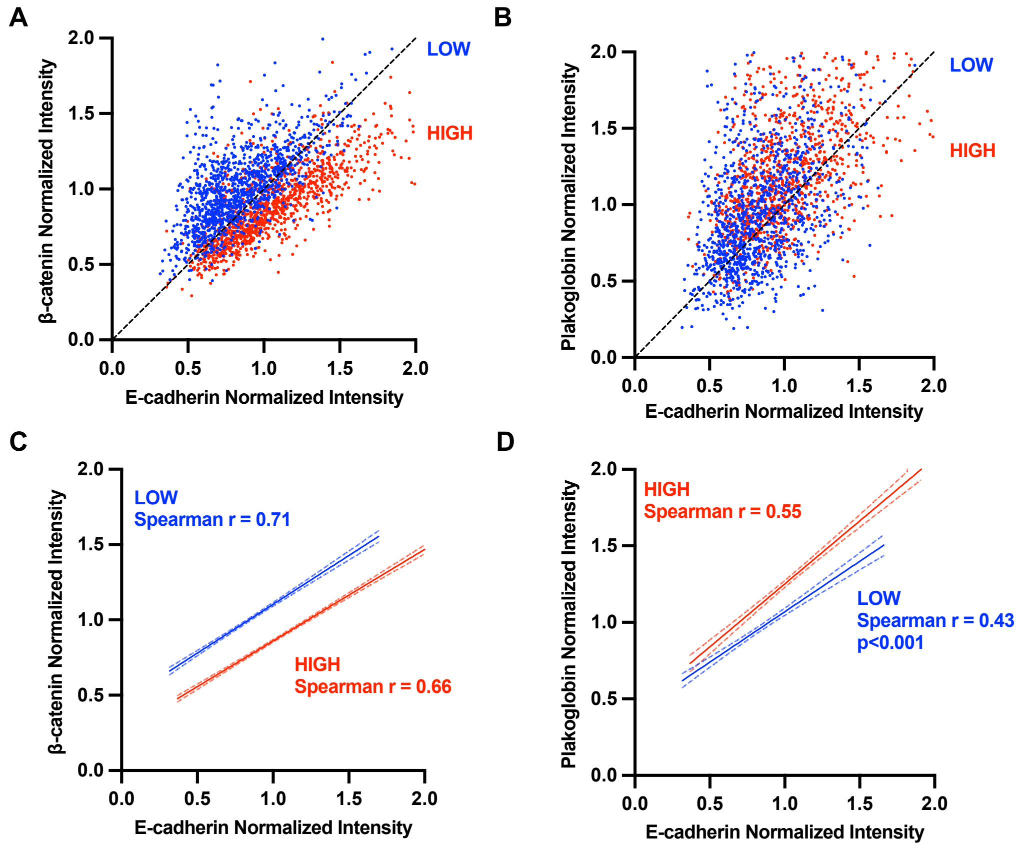


**Figure S6. Correlation of junctional E-cadherin, β-catenin, and plakoglobin.** Correlation plots from immunofluorescent data obtained from MDCK cells at low and high pHi co-stained with E-cadherin, β-catenin and plakoglobin. **(A-B)** Data shown are normalized fluorescence intensities of **(A)** β-catenin vs. E-cadherin and **(B)** plakoglobin vs. E-cadherin. From 3 biological replicates. All data were normalized to median values from control MDCK within each biological replicate. Dotted lines indicate identity (x=y). **(C-D)** Simple linear regressions shown with 95% confidence intervals. C is from data in A and D is from data in B. Shown in insets are Spearman correlation coefficients (r) for the data. P values indicated for statistically significantly different correlations when comparing across pHi.

A

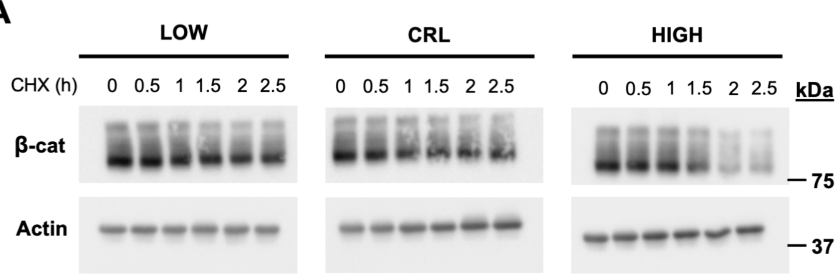


B

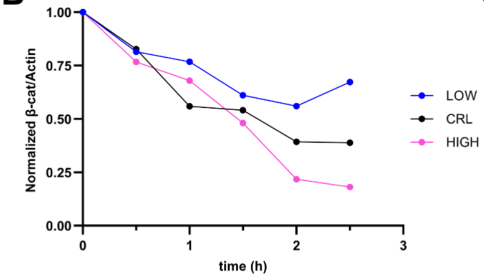


C

| Condition | Half-life (h) | $R^2$ |
|---|---|---|
| LOW | 2.79+0.03 | 0.75 |
| CRL | 1.58+0.03 | 0.96 |
| HIGH | 1.22+0.03 | 0.94 |

**Figure S7. Intracellular pH regulates the half-life of endogenous β-catenin. (A)** Representative immunoblots of β-catenin and actin. Cells were treated for pHi manipulation for 24 hours before being treated with cycloheximide (CHX) for the indicated amounts of time. **(B)** Quantification of immunoblots in (A) where β-catenin levels were normalized to actin. Ratios were normalized to t=0 h, plotted and fit with nonlinear regression. From one biological replicate. **(C)** Half-life values obtained using one-phase decay curve fitting for data in (B).

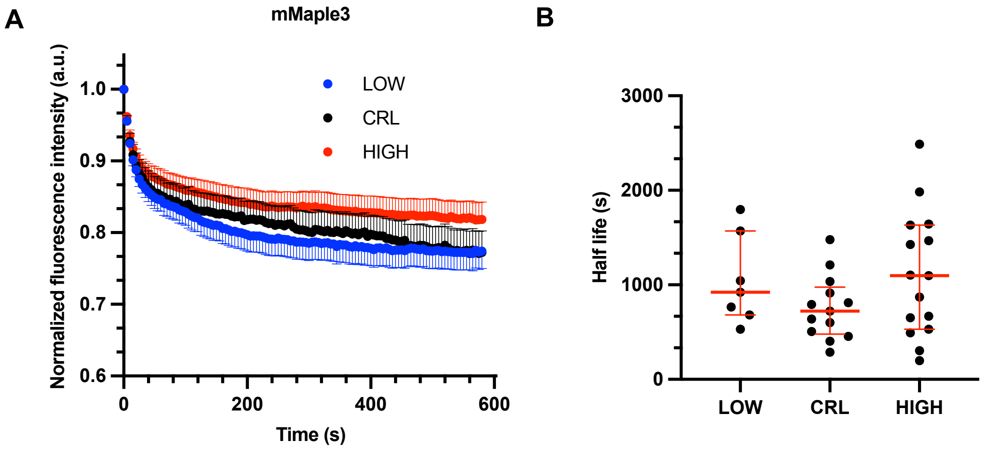


**Figure S8. mMaple3-β-catenin has pH independent degradation. (A)** Normalized fluorescence intensity traces from MDCK cells expressing mMaple3 under low, control, and high pHi conditions. Dots show mean cell traces, error bars indicate SEM. Reported half-life calculated from mean cell traces. LOW, n=14; CRL, n=9; HIGH, n=20 from 3 biological replicates. **(B)** Normalized fluorescence intensity traces from MDCK cells expressing mMaple3-β-catenin under low, control, and high pHi conditions. Dots show mean cell traces, error bars below (CRL) or above (HIGH, LOW) mean indicate SEM. Reported half-life calculated from mean cell traces. LOW, n=14; CRL, n=11; HIGH, n=18. **(C)** Single-cell half-life values were obtained by fitting individual single-cell traces from cells collected as in A and B. (median±IQR) Outliers were removed using the ROUT method (Q=1%). WT LOW, n=22; WT CRL, n=40; WT HIGH, n=29 from 6 biological replicates. H36R LOW, n=14; H36R CRL, n=11; H36R HIGH, n=18 from 3 biological replicates. Statistical significance in C was determined using Kruskal-Wallis test with Dunn's correction for multiple comparisons. *P<0.05, ****P<0.0001.
